# Modeling Hepatocellular Carcinoma and its microenvironment on a chip Hepatocellular carcinoma PDChip

**DOI:** 10.1101/2025.07.10.663127

**Authors:** Orsola Mocellin, Stephane Treillard, Abbie Robinson, Aleksandra Olczyk, Thomas Olivier, Chee P. Ng, Arthur Stok, Gilles van Tienderen, Monique M.A. Verstegen, Jeroen Heijmans, Dorota Kurek, Sebastian J. Trietsch, Henriëtte L. Lanz, Paul Vulto, Jos Joore, Karla Queiroz

## Abstract

Hepatocellular carcinoma (HCC) is the most common type of liver cancer. Its incidence is increasing and is closely related to advanced liver disease. Interactions in the HCC microenvironment between tumor cells and the associated stroma actively regulate tumor initiation, progression, metastasis, and therapy response. Effective drug development increasingly requires advanced models that can be utilized in the earliest stages of compound and target discovery. Here we report a phenotypic screen on an advanced HCC patient-derived chip (PDChip) model. The vascularized HCC PDChip models include relevant cellular players of the HCC microenvironment. We assessed the effect of 28 treatment conditions on a panel of 8 primary HCC tumors and 2 cell lines. Approximately 1200 HCC PDchips were grown under perfusion flow, exposed to treatments, and subsequently assessed for viability, tumor-associated vasculature responses and chemokine and cytokine changes.

Although the SoC therapeutics sorafenib and lenvatinib reduced culture viability and produced profound changes in the organization of the vascular beds, they did not affect the tumor cell population in these cultures. Atorvastatin, a 3-hydroxy-3-methylglutaryl-coenzyme A (HMG-CoA) reductase inhibitor, reduced tumor viability but did not affect vascular bed organization. Sorafenib, lenvatinib and atorvastatin also affected chemokine and cytokine release. Tocilizumab, galunisertib, and vactosertib decreased the level of IL6, a relevant prognostic marker for HCC, while IL6 was increased by halofuginone.

In conclusion, HCC PDChip models enabled a detailed evaluation of drug-induced responses in the tumor and associated microenvironment, highlighting their importance in preclinical research for understanding diseases and developing new drugs.

## Introduction

Hepatocellular carcinoma (HCC) is the third most common cause of cancer-related deaths worldwide and is a result of the malignant transformation of hepatocytes. HCC development is associated with viral disease, obesity, diabetes, and metabolic dysfunction-associated steatohepatitis (MASH), with approximately 90% of HCC developing on a background of cirrhosis ^12^. Until recently, sorafenib was the only first-line therapy approved by the Food and Drug Administration (FDA) for the treatment of advanced HCC ^3^. Currently, several multikinase inhibitors such as first-line treatment lenvatinib, and second-line treatments regorafenib, cabozantinib, and an anti-VEGFR2 antibody, ramucirumab, have also received FDA approval for advanced HCC^4^. However, the median overall survival for patients undergoing treatment remains under 15 months^5 4^. More recently, the FDA has approved nivolumab and pembrolizumab, and the combinations of nivolumab/ipilimumab, atezolizumab/bevacizumab, and tremelimumab/durvalumab, as second-line for sorafenib-pretreated HCC^6^. Nonetheless, tumor response rates for immune checkpoint inhibitors (ICIs) and combination therapies are reported to be only in the 8-20% range ^7^.Thus, a large unmet medical need remains for novel and effective (immuno)therapeutic strategies ^8^. Overall, HCC remains a highly lethal malignancy for the 40% of affected patients diagnosed with advanced-stage disease.

The tumor microenvironment (TME) of HCC is complex and has been shown to play a crucial role in HCC disease progression and response to treatment. Among relevant elements of the HCC TME are endothelial cells, stromal cells including liver stellate cells and cancer-associated fibroblasts (CAFs), and the tumor immune infiltrate. Advances in the understanding of the HCC TME indicate that these components potentially regulate each other and together influence extracellular matrix (ECM) remodeling, metastasis, cancer stemness, and therapeutic resistance ^9^. Commonly used HCC *in vitro* models such as cell lines, HCC-derived organoids, spheroids, and other 3D models, currently lack immune and supporting cells, only partially recapitulating disease complexity. Another important aspect, often overlooked by many *in vitro* platforms, is the ability to cultivate samples from different HCC patients thus assessing patient diversity and patient-specific responses^10^.

Given the limitations of HCC models, Organ-on-a-Chip (OoC) technology will potentially contribute to narrowing the gap between the relevance of HCC *in vitro* models employed in preclinical studies and *in vivo* biology as it allows for the inclusion of several key TME parameters such as the presence of vasculature, perfusion, supporting cell types, and the resulting cellular interactions and tissue organization.

On-chip systems have been used to model various cancers, including breast ^11^, pancreatic ^12^ ^13^ ^14^, colorectal, and lung^15^,providing advanced platforms for studying tumor cell dynamics and drug responses. Liver-focused on-chip systems have successfully simulated healthy and diseased states, replicating key features of liver tissue organization and metabolism. HCC-specific systems incorporating the HepG2 cell line have been used to model drug delivery ^16^ and bone metastasis ^17^. More recently, patient-derived HCC on-chip has been employed to study the impact of oxygen tension on tumor heterogeneity. This system has also been engrafted in mice, facilitating the growth of engrafted HCC tissue and generating multi-spot chip PDX models^18^.

Here, we advance the field of tumor modeling by combining a scalable microfluidic platform, the OrganoPlate, with HCC patient-derived tissue to simulate the cellular complexity of HCC and HCC patient heterogeneity *in vitro.* This comprehensive model setup, consisting of dissociated tumor tissue from HCC patients and HCC cell lines, combined with CAFs and vasculature, resulted in the generation of HCC on chip cultures able to produce the biomarker alpha-fetoprotein (AFP), and several chemokines and cytokines including IL6 and CCL2. HCC patient-derived chips (PDChips) were generated with samples from 8 different donors and 2 cell lines, and used to screen 28 different treatment conditions. The drug panel was composed of HCC SoC drugs, and a selection of drugs able to target several pathways in tumor cells as well as in the supporting cells, including endothelial cells and CAFs. Subsequently, all treatment responses were phenotypically characterized by assessing a combination of cell viability, vascular morphology and chemokine and cytokine production.

With this study, we demonstrate that the TME can be effectively modeled in a chip format, encompassing a range of cell types such as tumor and supporting cells, as well as important ECM components. Yielding robust cultures that can be screened across multiple donors and therapies. PDChip culture conditions facilitate relevant cellular interactions and organization, recapitulating disease complexity and patient diversity, allowing further insight into the targeting of HCC and its microenvironment. We believe that a broader implementation of tumor PDChip models will advance the development of novel and effective therapeutic alternatives.

## Results

### HCC PDChip models

To improve selection and evaluation of treatments targeting HCC, we developed an HCC PDChip model incorporating patient-derived tumor cells and components of the tumor microenvironment (TME) (Figure 1A). This comprehensive model setup includes primary dissociated HCC cells or HCC cell lines, HCC-derived CAFs, and endothelial cells. HCC tumors (stages I and II), Huh7 and HLE cell lines used to generate the HCC PDChip models are described in Table 1. Cultures were generated in the OrganoPlate Graft platform, containing 64 chips in each robot-compatible plate. Figure 1A shows the composition of the HCC PDChip model, the culture setup, and the readouts used in this study. HCC models were generated manually, whereas drug exposure, supernatant collection, addition of reagents, and culture fixation were automated and performed using a Biomek i5 pipetting robot (Beckman Coulter) and a MultiFlo FX non-contact dispenser (BioTek).

**Figure 1.**
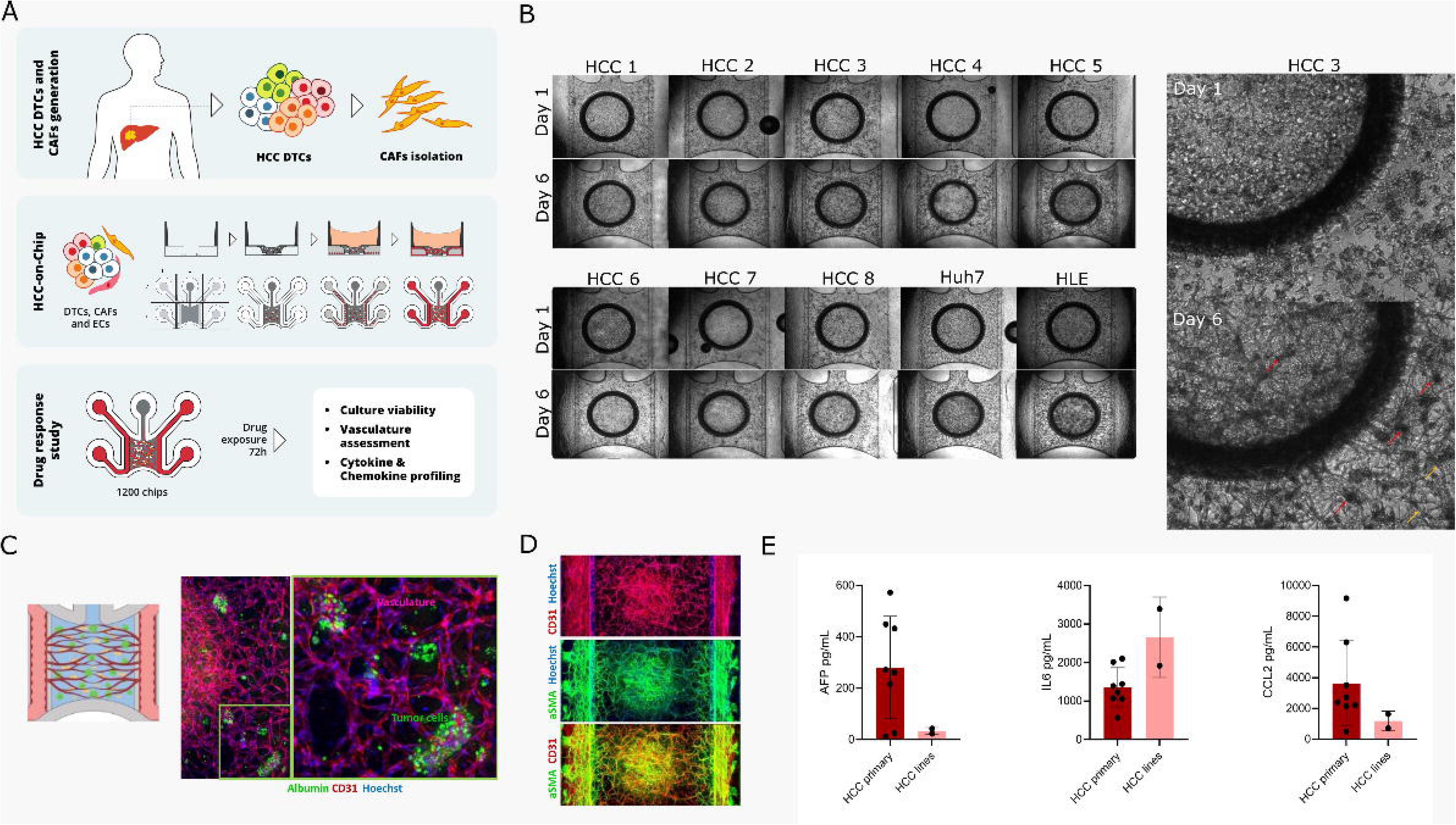
Generation and characterization of HCC PDChips. A. Schematic representation of HCC tissue processing and PDChips generation using the OrganoPlate Graft. DTCs, CAFs, and HUVEC were mixed in fibrin and dispensed into the chamber of the OrganoPlate Graft microfluidic platform where they were cultured for 6 days, followed by exposure to a selection compounds for 3 additional days. To assess compound responses, we evaluated cell viability, vascular network organization and chemokine and cytokine levels. B. Phase contrast images of tumor compartment and lining vasculature on day 1 and 6 of HCC PDChips, these cultures were generated using samples from 8 primary HCC tumors and 2 HCC cell lines. B. also shows the bottom right quadrant of the tumor compartment of a HCC3 PDChip, images show that cells are poorly organized on day 1, while on day 6 culture shows tumor aggregates and an organized vascular network. C. Confocal microscopy of HCC PDChip shows the presence of tumor cells (albumin, in green) and vasculature (VE-Cadherin, in red). D. Confocal microscopy of a HCC PDChip shows the presence of CAFs and vascular network, vasculature is immunostained for CD31 (red), CAFs are immunostained for aSMA (green) and nuclei are labeled with Hoechst (blue). Overlay image shows that CAFs seem to line vascular structures. E. shows the presence of AFP, IL6 and CCL2 (ng/mL) in HCC patient-derived and HCC cell lines models, HCC patients (N=8) and HCC cell lines (N=2; Huh7 and HLE). AFP, IL6 and CCL2 were measured in the supernatant collected from DMSO controls for all donors and cell lines on day 9.

**Table 1.**
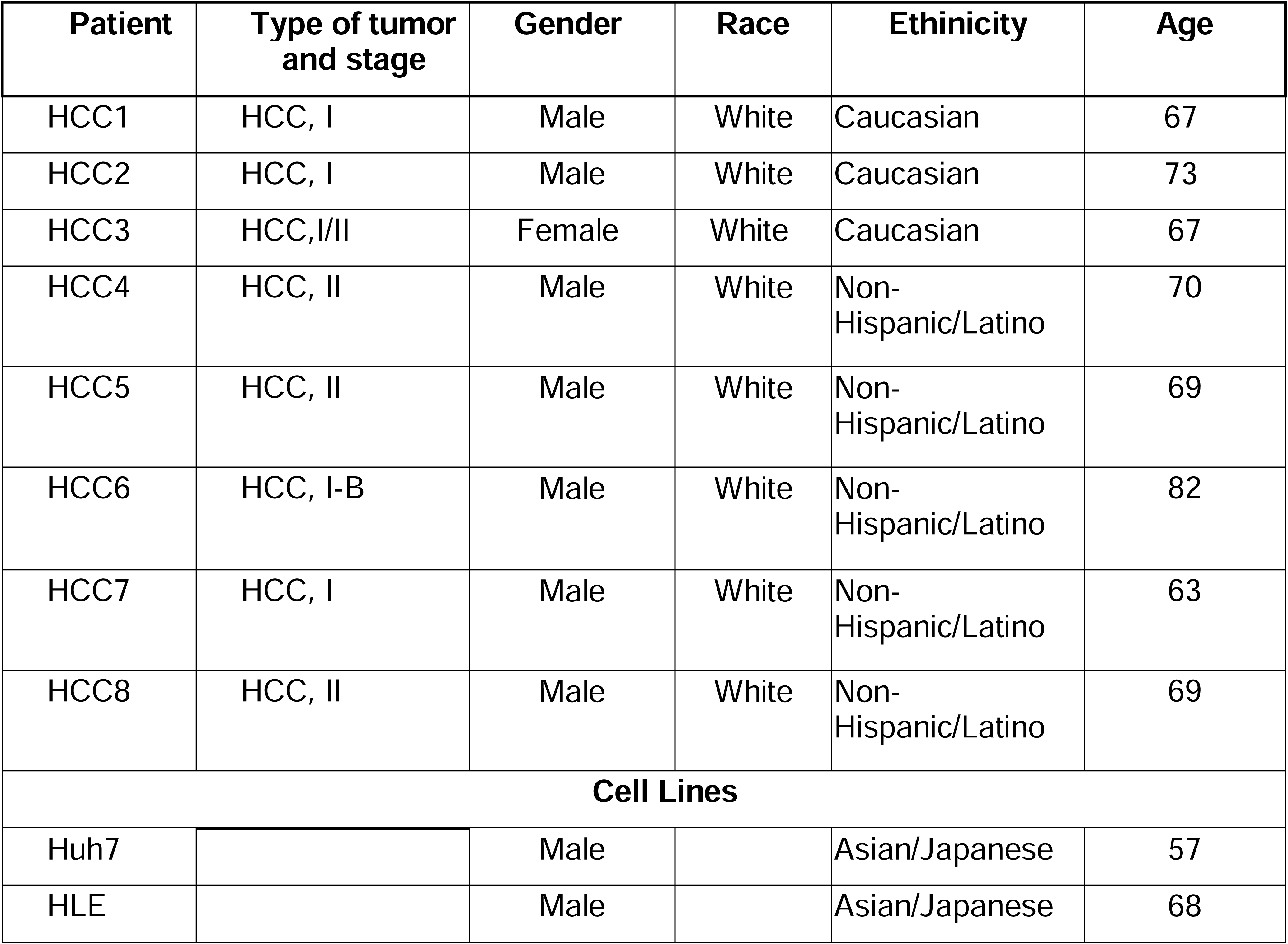
HCC patients information.

Dissociated primary HCC cells or cell lines, CAFs and HUVECs were resuspended in fibrin, loaded into a chip, and allowed to self-organize generating the tumor compartment (Figure 1A) (Bonanini et al., 2025).Side channels were loaded with endothelial cells, forming the vascular compartment (Figure 1A). HCC cultures developed well-organized vascular structures over time (Figure 1B and C). HCC tumor aggregates were visualized on phase contrast images on day 6 (Figure 1B). In addition, different cellular compartments were identified through immunostaining (Figure 1C). Interaction between tumor cell aggregates (albumin^+^) and vasculature (VE-Cadherin^+^) could be observed (Figure 1C). CAFs (α-SMA^+^) were observed lining the tumor vasculature (CD31^+^) as well as side endothelial tubules (perfusion compartment) (Figure 1D). Most HCC patient-derived cultures (6/8) presented levels of the HCC biomarker AFP >200ng/ml, and relevant secreted mediators such as IL6 and CCL2 were also present in these cultures. Very low levels of AFP were detected in the Huh7 and HLE cell line cultures.

HCC PDChips showed organized vasculature, and the presence of CAFs and tumor cells. AFP, IL6 and CCL2 levels in the supernatant differ across HCC patients. Next, cultures were exposed to several treatment conditions from day 6 to day 9.

### HCC PDChips viability in response to a drug panel

HCC PDChips were exposed to a panel of 28 treatment conditions (Table 2), including the HCC SoC treatments sorafenib and lenvatinib, and a selection of single and combination treatments (Table 2). Cultures were exposed on day 6 for 72 hours, after which supernatant was collected (Luminex analyses), cell viability evaluated (Alamar blue assay), and cultures were fixated for immunostaining.

**Table 2.**
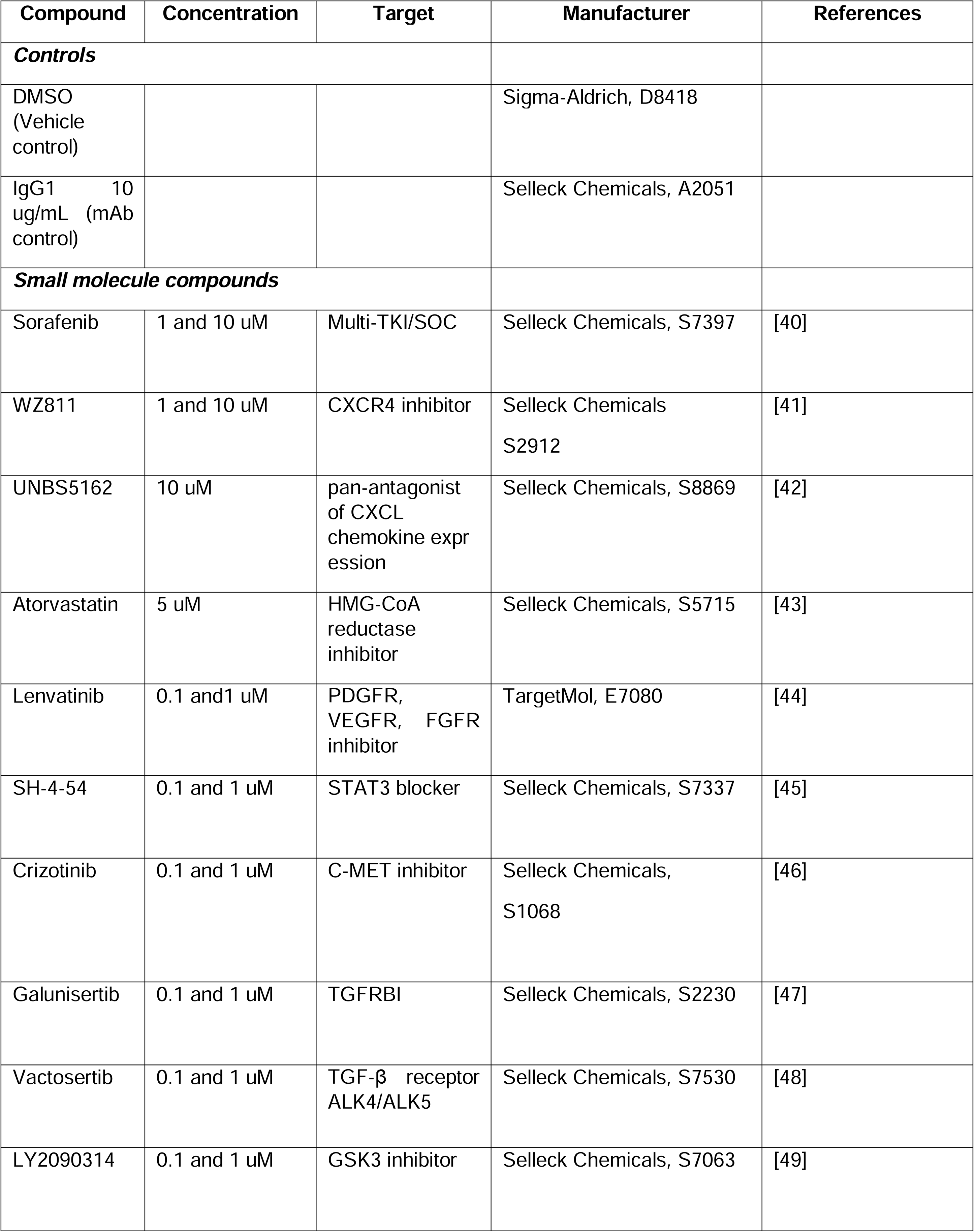
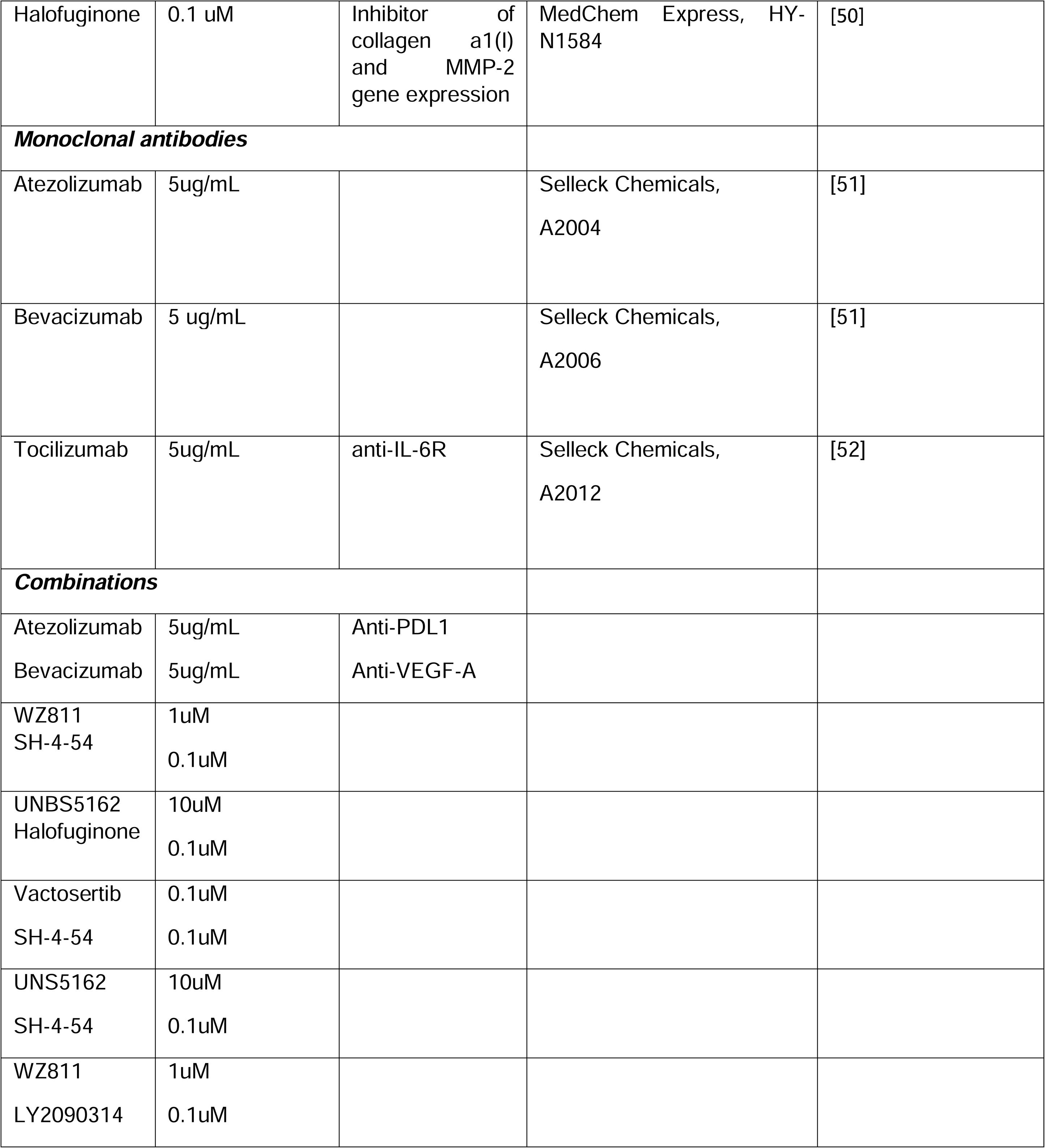
Compounds selection.

Viability assessment in the tumor and vascular compartments showed high reproducibility, indicating a comparable cell number and response between replicates (r=0.8236) (Figure 2A). Viability of DMSO controls was assessed to benchmark the distribution of the data per plate. To allow data analyses across different batches/plates, data from each plate was normalized to an average of the DMSO control response. In Figure 2B, DMSO control normalized data (normalized to average of DMSO control) is shown.

**Figure 2.**
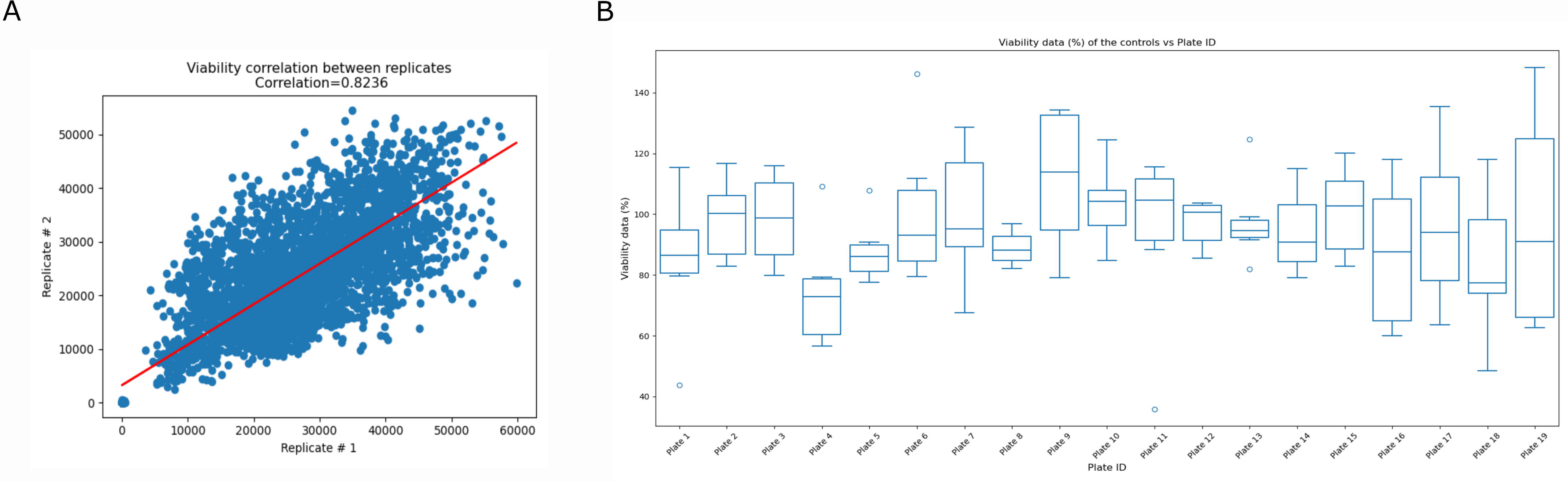
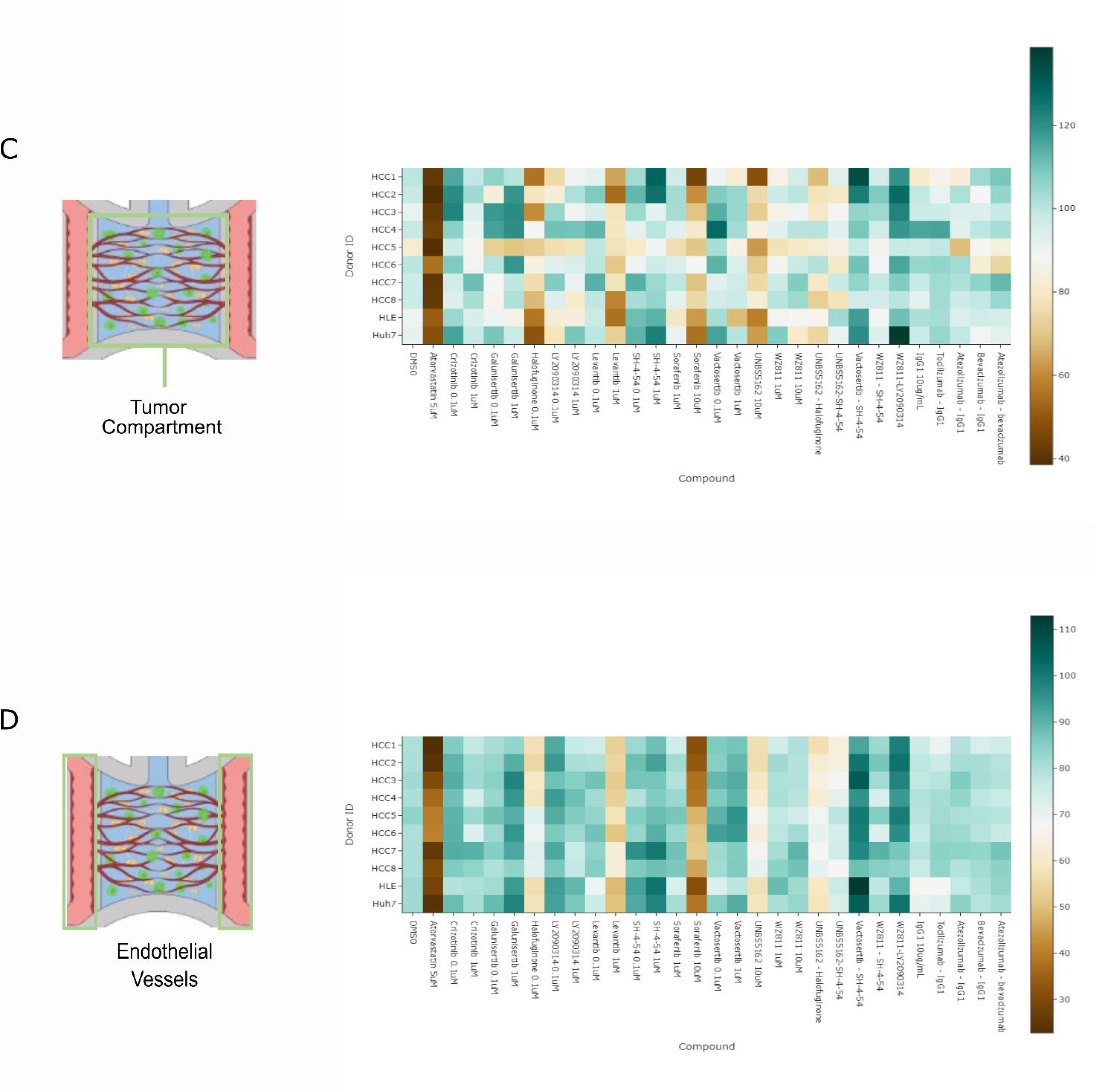
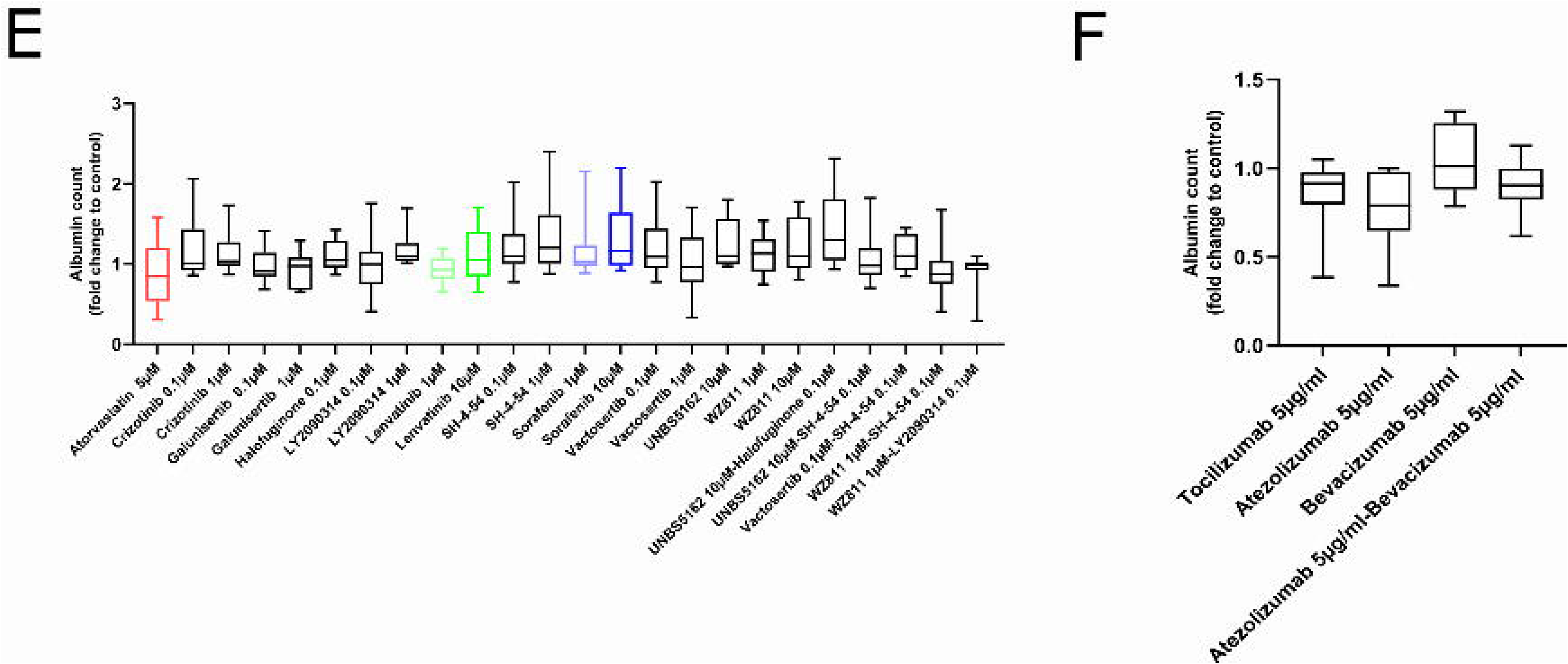
HCC PDChip model viability in response to CAF targeting and SoC. A. Scatterplot shows replicates reproducibility (r=0.8236) of PDChips viability results. DMSO control viability distribution across plates is shown in B. shows DMSO control viability percentage (values normalized to average of DMSO control viability/plate). C. Heatmap showing viability after treatment of the tumor compartment of HCC models (n=4-6). D. Shows viability after treatment of the vascular compartment of HCC cultures. Heatmaps represent viability percentages (normalized to average of all DMSO controls). E. and F. shows the quantification of albumin positive cells (tumor cells) in HCC PDChips in response to drug panel.

Viability of the tumor compartment decreased in response to sorafenib 10µM (71%), lenvatinib 1µM (80%), atorvastatin 5µM (47%), halofuginone 0,1µM (72%) and UNBS5162 10µM (72%) (Figure 2C). These drugs also decreased the viability of the vascular compartment (Figure 2D). After the Alamar blue viability assay, HCC PDChips were fixated and immunostaining for albumin, and CD31 was performed. Cultures were imaged on an ImageXPress Micro Confocal XLS (Molecular Devices) and images were analyzed using INCarta software (Molecular Devices). Tumor cell population in primary HCC-derived cultures showed a consistent distribution of albumin+ cells, compared to nearly absent levels in Huh7 cultures and a very inconsistent distribution in HLE cultures. Albumin+ cell population (tumor cells) was not significantly decreased in HCC patient-derived cultures in response to any of the treatment conditions (Figure 2E and F), suggesting that targeting of the TME or tumor cells by our drug panel did not result in HCC cell death.

Combined, these data provide further insight into how the culture, the vasculature, and the tumor cell population respond to compounds and compound combinations, including SoC treatments. The effect of these compounds on the organization of the tumor-associated vasculature was further investigated.

### Responses of tumor-associated vasculature to treatments

Angiogenesis plays a significant role in HCC carcinogenesis and progression (Moawad et al., 2020). Therefore, it was crucial to assess vascular responses of HCC PDChips to treatments. To study vascular responses to the drug panel, CD31 immunostaining was performed, and cultures were imaged by confocal microscopy (Supplementary Figure 1). Images of sorafenib and lenvatinib (HCC SoC) exposed cultures showed a clear effect of these drugs on the organization of the tumor-associated vasculature (Figure 3). Morphological changes promoted by these drugs are in line with their anti-angiogenic effects. Next, images from CD31 stained cultures (Supplementary Figure 1) were analyzed to assess parameters related to vasculature organization. First, a segmentation pipeline was applied to extract relevant parameters and evaluate compound-induced responses (Figure 3A). Several descriptors associated with vasculature organization were assessed (Table 3). tSNE and PCA plots including data extracted from analyses of vasculature images showed that sorafenib and lenvatinib induced similar changes on evaluated descriptors (Figure 3B and C). Assessment of tumor-associated vasculature confirmed morphological changes in line with sorafenib and lenvatinib anti-angiogenic effects. In Figure 3E-H selected parameters influenced by these drugs are shown (total vessel density, total vessel area, branching index and total vessel length). Additionally, LY2090314 0.1uM decreased vascular branching (Figure 3F). Despite the clear effect of Atorvastatin on cell viability of tumor and vascular compartments, this drug did not affect tumor-associated vasculature organization (Figure 3G).

**Figure 3.**
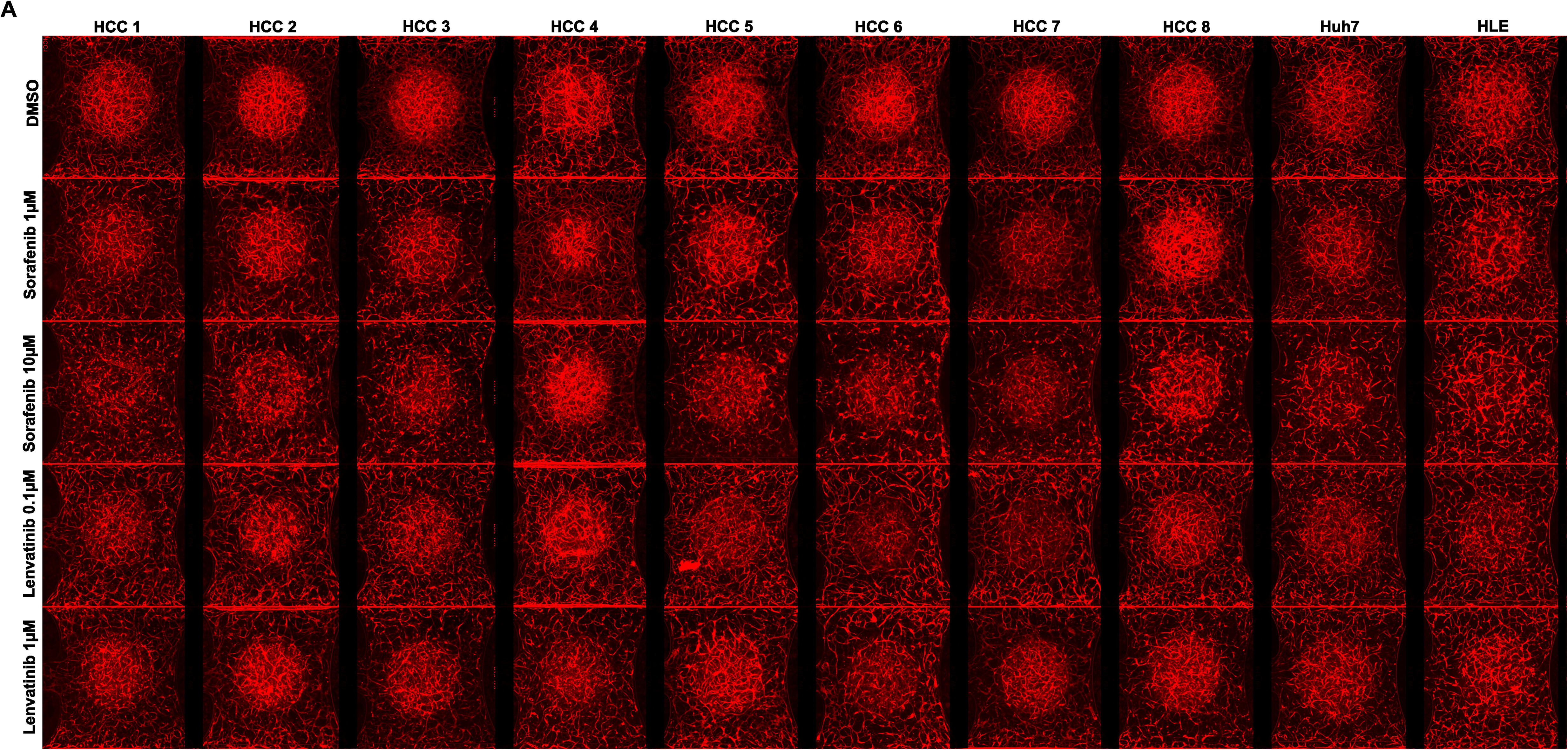
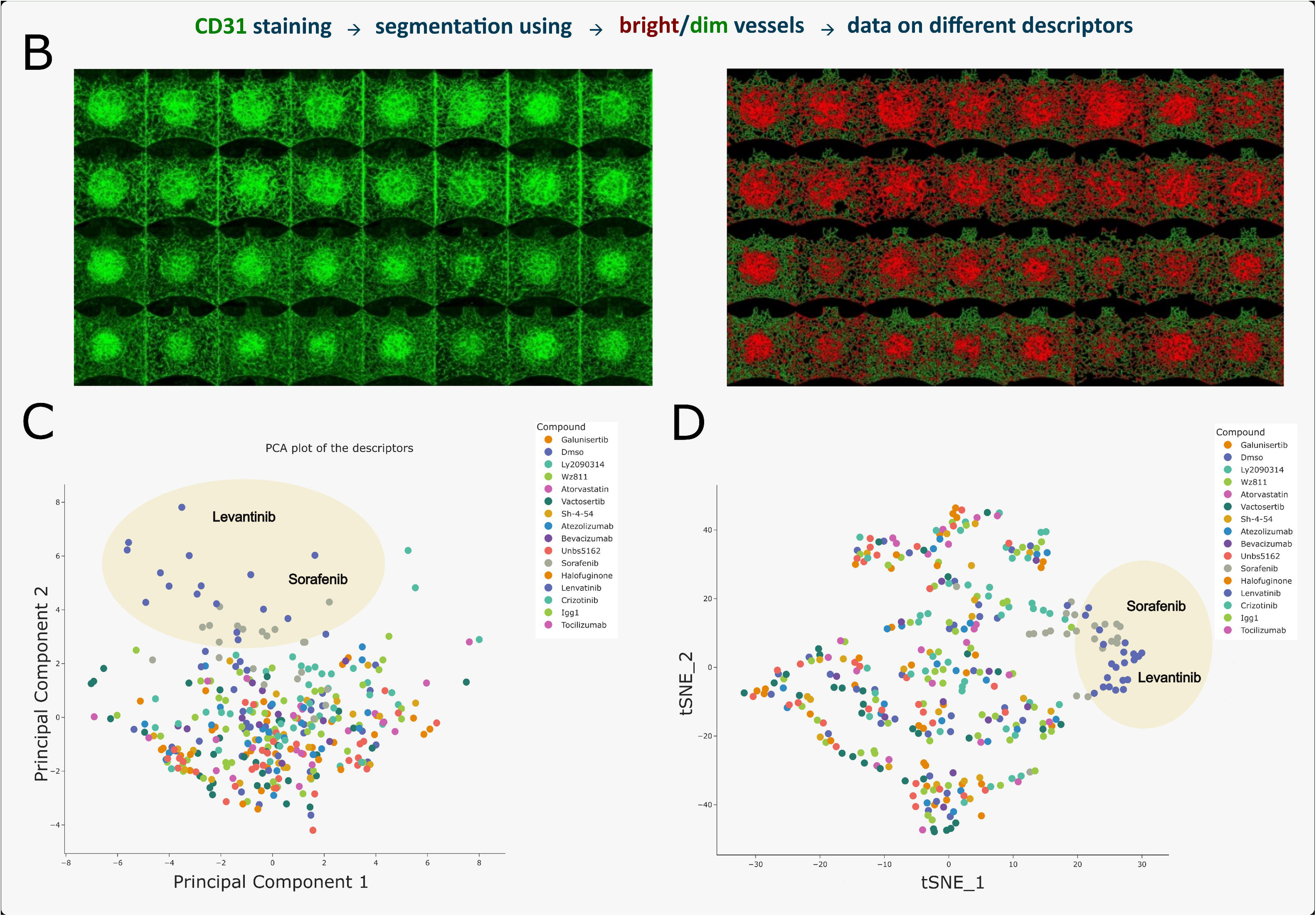
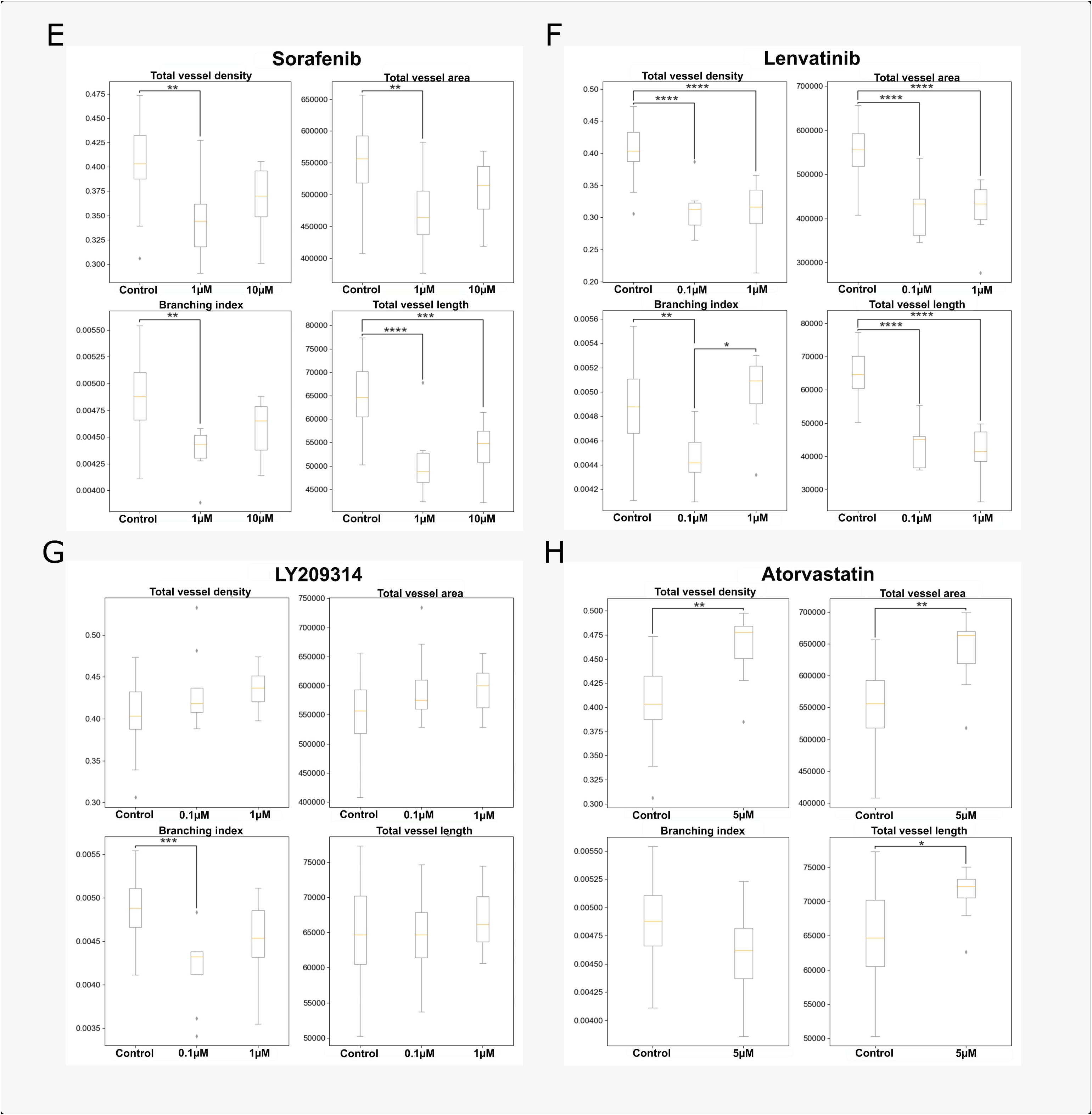
Treatment-induced vascular responses in HCC PDChips. **A.** HCC PDChip models associated-vasculature organization in response to sorafenib and lenvatinib. HCC cultures were immunostained for CD31 (red), and confocal imaged. B. Vasculature of HCC cultures was immunostained for CD31 and imaged using confocal microscopy (CD31 immunostaining for all cultures and treatment conditions can be found in Supplementary Figure 1) (n=30, for DMSO control; n= 10, for each treatment condition). Images were processed and 15 vascular organization-related descriptors were extracted. C. Unsupervised linear (PCA), and D. non-linear (t-SNE) reduction of the data show a clustering of sorafenib and senvatinib. E-H shows sorafenib, lenvatinib, LY2090314 and atorvastatin-induced changes in total vessel density, total vessel area, total vessel length and branching index. Graphs 4E-H include HCC patient-derived and HCC cell line culture results. Statistical significance was determined using either the Kruskal-Wallis test followed by Dunn’s post hoc test or ANOVA followed by Tukey’s post hoc test, depending on whether assumptions for parametric tests were met, with p-values adjusted using the Bonferroni correction. Adjusted p-values are expressed as follows: **** p<0.0001, *** p<0.001, ** p<0.01, * p<0.05.

**Table 3.**
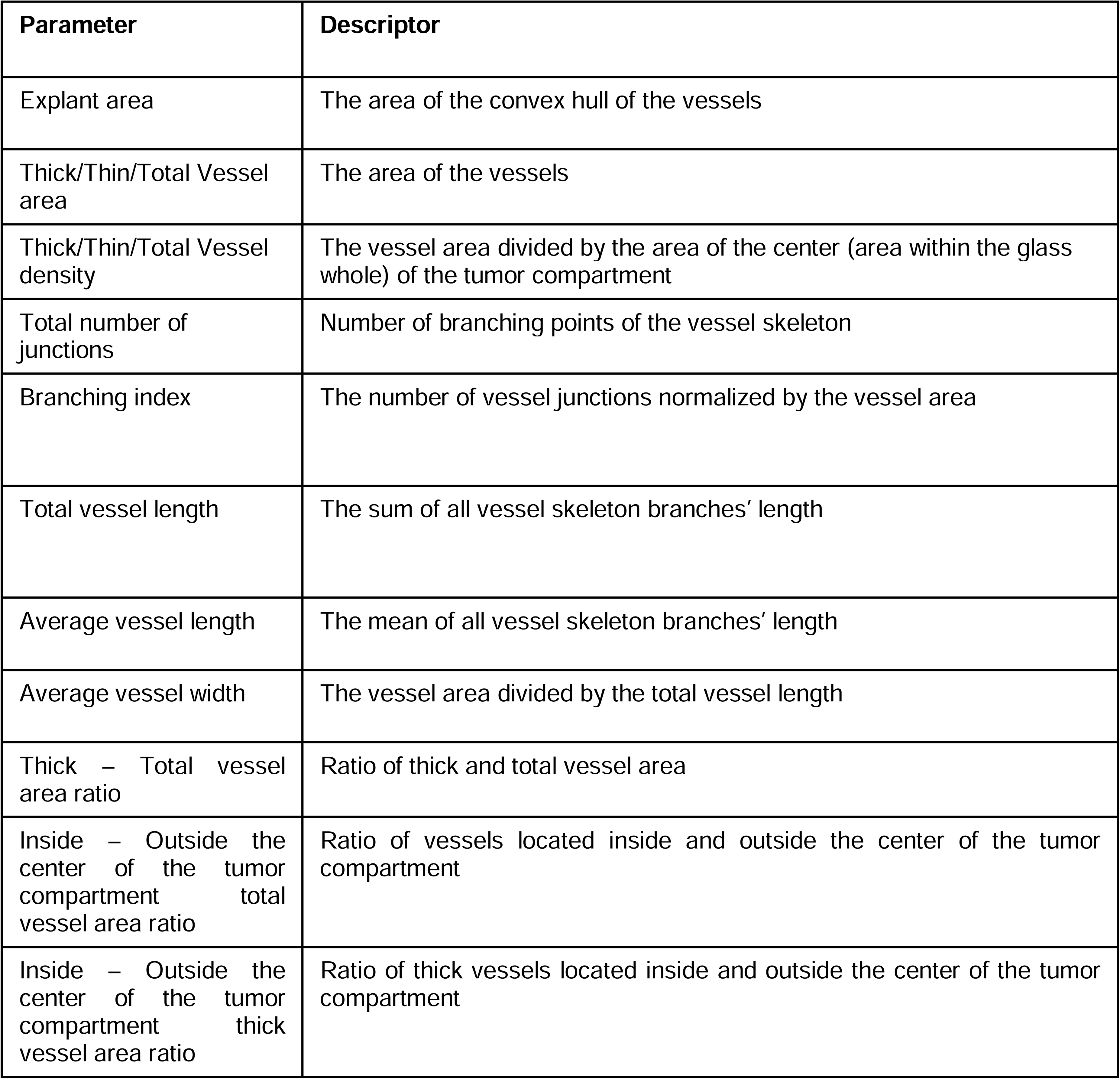
Parameters and descriptors used for phenotypical assessment of tumor-associated vascular network.

| Parameter | Descriptor |
| --- | --- |
| Explant area | The area of the convex hull of the vessels |
| Thick/Thin/Total Vessel area | The area of the vessels |
| Thick/Thin/Total Vessel density | The vessel area divided by the area of the center (area within the glass whole) of the tumor compartment |
| Total number of junctions | Number of branching points of the vessel skeleton |
| Branching index | The number of vessel junctions normalized by the vessel area |
| Total vessel length | The sum of all vessel skeleton branches' length |
| Average vessel length | The mean of all vessel skeleton branches' length |
| Average vessel width | The vessel area divided by the total vessel length |
| Thick – Total vessel area ratio | Ratio of thick and total vessel area |
| Inside – Outside the center of the tumor compartment total vessel area ratio | Ratio of vessels located inside and outside the center of the tumor compartment |
| Inside – Outside the center of the tumor compartment thick vessel area ratio | Ratio of thick vessels located inside and outside the center of the tumor compartment |

HCC PDChips exhibit a very organized vascular network, which is only globally affected by SoC, sorafenib and lenvatinib. Next, cytokines and chemokines levels in HCC PDChips in response to the drug panel were analyzed.

### Drug-induced immunomodulatory response in HCC PDChips

The HCC TME is highly complex, and the combinations of different cell types and resulting interactions have a significant influence on tumor progression, drug sensitivity, and the tumor immune landscape. Considering that supporting cells such as CAFs and endothelial cells are known to influence immune cell recruitment and function, we measured a panel of immunosuppressive (CXCL1, CXCL8, CXCL12, CCL2, CCL20, IL4, IL6) and immunostimulatory (IL21, TNF, CXCL10, CXCL11, CCL3 and CCL4) chemokines and cytokines, and the HCC biomarker AFP. Culture supernatant levels of these analytes were measured in response to the different treatments.

Supernatant analysis results show significant changes in multiple chemokines and cytokines (Figure 2 and 4) induced by treatments in HCC patient cultures. A heatmap visualization (Figure 4B) provides an overview of the effect of different treatments on chemokine and cytokine levels in HCC patient cultures. HCC biomarker AFP remained unaffected by treatments. Sorafenib exposure resulted in reduced levels of several chemokines and cytokines (CCL4, CXCL1, CXCL10, IL21, IL4 and TNFα) potentially promoting an immunosuppressive effect. A similar response was observed in the presence of lenvatinib, although CXCL12 was increased in response to lenvatinib 0.1uM. Atorvastatin induced a decrease in most tested analytes, this was likely attributable to its effect on culture viability.

**Figure 4.**
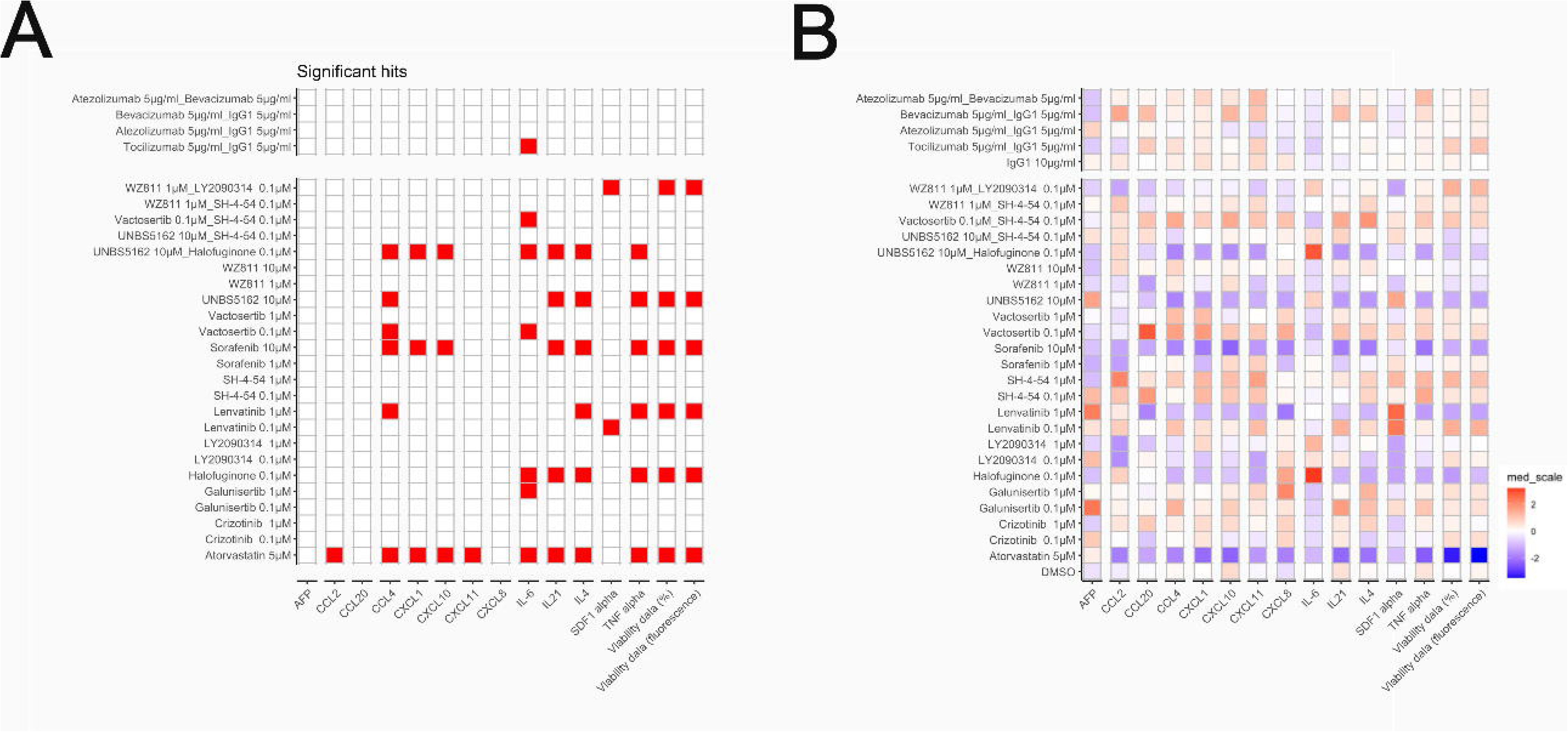
Immunomodulatory response in HCC PDChip models. Supernatant of HCC cultures was collected and analysed for expression of a panel of 13 cytokines and chemokines and IL6 by Luminex. A. Significant changes in cytokine and chemokine production in HCC PDChip models in response to treatments. B. Heatmap shows an overview of cytokine and chemokine responses of HCC cultures in response to treatments. (n=30, for DMSO control; n= 10, for treatment conditions). Statistical significance was determined using Wilcoxon tests, with p-values adjusted using the Benjamini-Hochberg correction. Adjusted p-values are expressed as follows: **** p<0.0001, *** p<0.001, ** p<0.01, * p<0.05.

IL6, which is a prognostic biomarker^20^ in HCC, was significantly affected by several treatments, showing reduced levels in response to tocilizumab, galunisertib and vactosertib, whereas halofuginone alone or in combination with UNBS5162 led to increased IL6 levels. Halofuginone also promoted a decrease in IL4, IL21 and TNFα.

These data suggest that targeting of the supporting cell compartment influences relevant immune mediators, and that the use of such compounds could potentially impact patient anti-tumor immune responses.

## Discussion

HCC remains challenging to treat, with a 5-year overall survival after diagnosis observed in only 18% of patients. This low survival rate is usually associated with late diagnosis, and limited treatment options^21^. Approved treatment options target different HCC tumor components, including the tumor, vascular, and immune compartments, indicating that the modulation of the TME is essential for effectively treating this tumor type ^22^. Equally relevant is the variability across patients, which underscores the need for patient-derived models and for tailored therapeutic management of specific patient subsets ^23,10^.

Several *in vivo* and *in vitro* models for HCC have been developed to understand HCC biology and drug sensitivities. While these models contribute in a relevant manner to the understanding of HCC biology, recapitulating HCC complexity and HCC patient diversity is still challenging and requires further efforts

Here we report on an HCC PDChip setup that brings us a step closer to creating more comprehensive HCC models that are able to capture relevant HCC cellular self-organization and interactions. HCC patient-derived dissociated tumor tissue were cultured in presence of endothelial cells and CAFs. These cultures were generated on a scalable OoC platform and maintained for 9 days. Cellular interactions within the microfluidic platform led to vascularized tumor constructs, with HCC aggregates being enveloped and traversed by a lumenized and interconnected vascular plexus, in association with CAFs. Biomarkers AFP and IL6, as well as CCL2 were present in the supernatant of these tumor constructs (Figure 1).

Vascularized HCC PDChips were exposed to a panel of treatment conditions that included SoC, single and combination treatments (Table 2). Drugs were selected for their ability to target tumor cells, endothelial cells and/or CAFs, and potentially modify cultures viability, organization as well as chemokine and cytokine landscape.

PDChips showed decreased viability in response to sorafenib 10µM, lenvatinib 1 µM, halofuginone 0.1 µM, UNBS5162 10 µM and atorvastatin 5 µM; while WZ811 1 µM in combination with LY2090314 0.1 µM increased viability of HCC cultures (Figure 2D). This effect remained significant when the whole group of HCC patient response was compared to controls (Figure 4A and B). Quantification of tumor cells (albumin+ cell population) in these cultures suggested that, on average, this population remained unaffected by tested treatments as well as SoC (sorafenib and lenvatinib). Response to SoC seems to align with clinical data, sorafenib and lenvatinib have been shown in multiple trials to prolong median survival and the time to progression. However, most responders exhibit stable disease rather than significant tumor regression in response to these drugs ^3^ ^24^. Atezolizumab plus bevacizumab did not affect HCC PDChip cultures viability, which could be a result of underrepresentation of immune cells in the DTC population or poor maintenance of this cell population under the culture conditions used.

Next, we phenotypically characterized the associated vasculature of treated HCC tumor constructs. Cultures exposed to SoC, i.e. sorafenib and lenvatinib, were characterized by disorganization of the vascular beds compared to the control condition (Figure 3). This response could potentially be associated with the decrease of tumor vascularity normally observed in response to these compounds in patients^22^.LY2090314, a GSK3a/b inhibitor, specifically affected tumor vasculature branching. As discussed previously, atorvastatin decreased HCC culture viability and did not result in abnormal vasculature. However, this drug did promote an increase in total vessel density, total vessel area, and total vessel length. Reported endothelial cell responses to atorvastatin are complex, dose- and context-dependent, and include effects such as anti-angiogenesis, cytoprotection, and promotion of vessel maturation^25^ ^26^ ^27^ ^28^.

Culture supernatant was profiled using a Luminex panel. Treatments did not influence AFP levels (Figure 4A). Lenvatinib and sorafenib in general negatively influenced chemokine and cytokine production (Figure 4A and B). A similar response to sorafenib has also been observed previously in explant cultures^29^. Interestingly, IL6, which is an important mediator in HCC associated with a poorer response to sorafenib, regorafenib as well as atezolizumab plus bevacizumab^30^ ^31^ ^32^, was decreased in HCC cultures supernatant in response to atorvastatin, galunisertib, vactosertib and tocilizumab. In contrast, halofuginone increased IL6 levels. In addition, this compound, known for its anti-fibrotic effects, decreased the levels of TNFα, IL4 and IL21. This was also observed when halofuginone was used in combination with UNBS5162. Halofuginone has demonstrated anti-fibrotic activity by reducing collagen I gene expression, regulating MMPs activity, increasing TIMP synthesis, and inhibiting TGFβ-driven signaling ^33^ ^34^. Additionally, halofuginone inhibits prolyl-tRNA synthetase, activating the amino acid starvation response (AAR) pathway, leading to increased ATF4 expression ^35^. ATF4 has been shown to link metabolic stress to increased IL6 expression ^36^, which may explain the observed rise in IL6 levels in HCC cultures treated with halofuginone.

While patient-derived models are invaluable for cancer research due to their clinical relevance, they come with several limitations. Tumor heterogeneity poses a significant challenge, as variations between patients can limit standardization. Additionally, the complexity and cost of developing and maintaining these models are substantial. Scalability is another challenge, making it difficult to use these models for high-throughput screening or large-scale studies. Despite these challenges, patient-derived models remain a crucial tool, offering insights that are often more clinically relevant than traditional cell lines or animal models. However, detailed analyses of these tumor models enabled by the combination of several techniques seem essential to understanding model responses to different molecules as well as leveraging on the unmatched capabilities of on chip systems to recreate complex cellular interactions in vitro.

Altogether, our model setup enables testing of tumor, CAF and vascular targeting molecules, potentially also allows for the understanding of the role of tumor-vasculature or tumor-CAF interactions in tumor cell behavior and drug responses. HCC PDChip reported here has the potential to provide a platform that includes intra- and interpatient variability with sufficient scalability and ease of use for industrial and clinical implementation.

## Materials and Methods

### Cells

Human umbilical vein endothelial cells (HUVEC, Lonza, C2519AS) were cultured in complete MV2, consisting of Endothelial Cell Growth Medium MV2 (PromoCell, C-22022) supplemented with 1% penicillin/streptomycin (Sigma Aldrich, P4333).

Cancer-associated fibroblasts (CAFs) from donor HCC3 were isolated in-house and cultured in a combination medium composed of 67% complete MV2 and 33% complete EMEM, consisting of EMEM (ATCC, 30-2003) supplemented with 10% fetal bovine serum (FBS, Gibco, 16140-071) and 1% penicillin and streptomycin solution (Sigma Aldrich, P4333).

Dissociated tissue cells (DTCs) from HCC tumor tissues HCC1 and HCC3 were obtained in collaboration with the Department of Surgery, Erasmus MC-University Medical Center, The Netherlands. Tumor biopsies collected during surgical removal of the tumor for curative intent, were kept on ice until use. METC approval (MEC2013-143) and written informed consent to use the biopsies for research purposes was provided by the patients. Samples were confirmed to be of tumor-origin with histopathological assessment. These were dissociated in-house following the protocol described below (Tissue dissociation in Supplemental information) and cryopreserved in freezing medium (90% FBS, 10% DMSO) until seeding. DTCs from donors HCC2, HCC4, HCC5, HCC6, HCC7, and HCC8 were purchased from Discovery Life Sciences and cryopreserved until seeding (Table 1). Informed consent was obtained from all donors by the tissue provider, and ethical oversight was managed by the tissue provider and MIMETAS.

HCC cells lines Huh7 (JCRB0403) and HLE (JCRB0404) were obtained from the Japanese Collection of Research Bioresources Cell Bank. These were cultured in DMEM supplemented with 10% fetal bovine serum (FBS,Gibco 16140-071) and 1% of penicillin and streptomycin solution (Sigma Aldrich, P4333).

### Tissue dissociation

Fresh HCC tissue was dissociated as follows: the tissue was washed with Ad+++ (advanced DMEM/F12, Thermo Fisher Scientific,12634010) supplemented with 1%penicillin and streptomycin and cut into 3-5 mm pieces. The pieces were then collected in DBSA, DMEM supplemented with 1% bovine serum albumin (BSA, Sigma Aldrich, A2153) and centrifuged for 10 minutes at 300g. The supernatant was discarded, and the pellet was resuspended in digestion medium (Ad+++ supplemented with 2.5mg/mL collagenase IV (STEMCELL Technologies, #07427) and incubated at 37°C with vigorous shaking. After 15 minutes, DNase (Sigma, D5025) was added to the digestion medium and incubated for 30 minutes at 37°C with vigorous shaking. The digestion was stopped by addition of complete EMEM and the cell suspension was passed through a 70µm cell strainer. The resulting single-cell suspension was centrifuged for 10 minutes at 400g, the supernatant was discarded, and the cell pellet was resuspended in ammonium chloride solution (STEMCELL Technologies,07800) and incubated for 2 minutes to lyse any remaining red blood cells. The lysis was stopped by addition of DBSA, then cells were counted and either cryopreserved in freezing medium or cultured for fibroblast isolation.

### Fibroblast isolation

Cancer-associated fibroblasts (CAF) were isolated from donor HCC3 dissociated tissue cells (DTC). Following HCC tissue dissociation, cells were cultured in a T25 flask in 67% complete MV2 and 33% complete EMEM, consisting of EMEM (ATCC, 30-2003) supplemented with 10% fetal bovine serum (FBS, Gibco, 16140-071) and 1%penicillin/streptomycin solution (Sigma Aldrich, P4333) until formation of fibroblast colonies, as established by observation under light microscopy based on morphology. Cells were then passaged and cultured until passage 4, when cell population consisted mostly of fibroblasts as determined by morphology and α-SMA expression (detected by immunofluorescence).

### Generation of HCC PDChips

Cells were seeded into an OrganoPlate^®^ Graft (MIMETAS B.V, 6401-400-B). Seeding was performed as follows; HUVEC and CAF were cultured for one passage prior to seeding (seeded at passage 6 and 5, respectively). On the day of seeding, cells were detached using Trypsin/EDTA (Lonza, CC-5012), neutralized using complete MV2, centrifuged for 5 minutes at 300g and resuspended in complete MV2 to the desired concentration. DTC were thawed in complete MV2, centrifuged for 5 minutes at 300g and resuspended in complete MV2 to the desired concentration. Thrombin (Enzyme Research Laboratories, HT 1002A) was diluted in complete MV2 to a concentration of 1U/mL. Cells, thrombin, and fibrinogen (Enzyme Research Laboratories, FIB1) were kept on ice through the seeding process. The seeding mixture was prepared in the ratios described below by combining all cell types first, followed by addition of fibrinogen and thrombin immediately prior to seeding. Seeding mixture (1.35μL) was dispensed in the graft chamber and allowed to polymerize at room temperature for 10 minutes. Following polymerization, 1μL HUVEC suspension (10 000 cells/μL in complete MV2) was added to each perfusion channel inlet and allowed to fill the perfusion channels before dispensing 50μL complete MV2 to the perfusion channel inlets and to the graft chamber. The plate was incubated static at 37°C for approximately 2 hours to allow for HUVEC attachment in the perfusion channels. 50μL complete MV2 was then dispensed in the perfusion outlets and the plate was incubated at 37°C on the OrganoFlow at 14° at an 8-minute interval. On the next day, all medium was removed from the plate and replaced with 50uL/well complete ECGM2 (Endothelial Cell Growth Medium 2, PromoCell C-22011, supplemented with 1% p/s). Complete ECGM2 was refreshed in all wells every 48 hours until compound exposure.

### Compound exposure

The drug panel (Table 2) was composed by HCC SoC as well as a selection of compounds able to target several signaling pathways in tumor cells as well as the supporting endothelial cells and CAFs. Compound exposure was performed on day 6 of culture, when culture medium was removed and replaced with complete ECGM2 containing treatment compounds or controls according to the table below (Table 2). Each treatment condition and monoclonal antibody control was assigned to two chips in each plate (four chips per donor), while our vehicle control was assigned to six chips in each plate (twelve chips per donor). Compound addition to the plate was performed semi-automatically using a pipetting robot (Biomek i5; Beckman Coulter) to mix and dispense the reagents. Compound exposure was maintained for 72 hours, until the end of the culture (day 9).

### Alamar blue viability assay

On culture day 9, cell supernatant was collected from each well using a pipetting robot (Biomek i5; Beckman Coulter) and stored at −80°C until cytokine analysis. Following supernatant collection, AlamarBlue cell viability reagent (Thermo Fisher Scientific, DAL1100) diluted 1:10 in culture medium was added to each well using a non-contact dispenser (MultiFlo FX; BioTek) and incubated at 37°C for 2 hours on the OrganoFlow^®^ rocker (MIMETAS B.V., MI-OFPR-L) at 14° at an 8 minute interval. Following incubation, fluorescence was quantified at 544/590nm using the Spark Cyto plate reader (Tecan Life Sciences).

### Immunohistochemistry

After Alamar blue assay cultures were fixed as follows; all medium was removed from each well, replaced with fixation solution 3.7% formaldehyde (Thermo Fisher, 033314.K2) in HBSS with calcium and magnesium (Thermo Scientific, 14025092), and incubated at room temperature for 15 minutes. Fixation solution was removed, and each well was washed three times with phosphate buffered saline (PBS). Following the third wash, 50uL PBS were added to each well and the plate was sealed using Parafilm and stored at 4°C until immunofluorescent staining. For the immunofluorescent staining of endothelial, tumor, and fibroblast markers, plates were permeabilized and blocked for 2 hours with blocking buffer (1% Triton X-100 +3% BSA in PBS), followed by primary antibody incubation for Albumin-FITC (A80-229F, Bethyl), VE-Cadherin (Ab33168, Abcam), CD31 (M0823, Dako) and α-SMA (A2547, Sigma). Nuclei were stained with Hoechst (H3570, Thermo Fisher). Primary antibodies were diluted in antibody buffer (0,3 % Triton X-100 3% BSA in PBS) overnight. Primary antibodies were washed three times with washing buffer (0.3% tritonX-100 in PBS) and incubated with Donkey anti-rabbit 647 (A31573, Thermo Fisher) or Goat anti-mouse 647 (A21236, Thermo Fisher) secondary antibodies in antibody buffer overnight. Secondary antibodies were washed three times with washing buffer and once with PBS. Following the washing steps, PBS was added to all wells and the plate was imaged using the ImageXPress XLS Micro Confocal (Molecular Devices).

### Luminex analysis

Medium from culture day 9 was collected for cytokine and chemokine analysis. The samples were analyzed for quantification of AFP, CCL2, CCL3, CCL4, CCL20, CXCL1, CXCL10, CXCL11, IL-4, IL-8, IL-21, CXCL12, TNFα with the Human ProcartaPlex Mix&Match 13-plex (ThermoFisher Scientific) and for quantification of IL-6 with ProcartaPlex Human IL-6 (ThermoFisher Scientific). The assay was carried out following the protocol provided by the manufacturer. Raw data was processed using the ProcartaPlex Analysis App (ThermoFisher Scientific).

### Vascular bed analysis

To assess phenotypical changes in the vascular network of HCC cultures, images were processed using FIJI, and segmented using a trainable image processing plugin Labkit ^37^. Vascular network images (visualized through CD31 immunostaining) were preprocessed, and background signal was removed using the rolling ball background subtraction algorithm ^38^. This step removed unwanted artifacts from the vessel images. Next, images were fed to a classifier trained with the Labkit plugin, resulting in a three classes segmentation: background, thin vessels and thick vessels (Figure 1B). After being segmented, the images were postprocessed, where signal outside of the area of interest (tumor compartment) and artifacts smaller than a certain threshold were removed, this improved the quality of the segmentation. The cleaned segmented network was then skeletonized to extract vessel descriptors further described in Table 3. Further data analysis was performed using Python The Python packages Pandas (Mckinney, 2010), NumPy ^39^ and Scikit-learn (Pedregosa et al., 2011) were used for data processing and Matplotlib was used for visualizations.

### Data analysis

Data was processed in Excel and graphs were generated in GraphPad, R studio and Python. Raw Luminex data was processed using the ProcartaPlex Analysis App (ThermoFisher Scientific), data analysis and visualization were performed using R version 4.4.2 and RStudio version 2024.12.0 using the tidyverse package set. Normality was assessed using Shapiro-Wilk test. Statistical analyses were performed using pairwise Wilcoxon rank-sum test with Benjamini-Hochberg correction for multiple testing. Vascular beds analyses were performed using CD31 images, these were segmented using a Labkit based segmentation model in FIJI. Albumin expression was quantified using INCarta image analysis software (Molecular Devices, version 2.1). A U-Net CNN model based on ResNet was trained using the SINAP module. The model accurately segmented the Albumin staining under a range of different expression levels. The total area expressed per chip was extracted, linked with experimental treatment metadata for comparison through bar charts. For the comparison of viability, tumor cell population, vasculature and morphology independent analyses were conducted. One-Way ANOVA followed by Tukey’s HSD post-hoc test was used for gaussian data, or Kruskal-Wallis followed by Dunn’s post-hoc test for non-gaussian data was used. All p-values were adjusted for multiple comparisons using Bonferroni correction, with adjusted values (q-values) ≤ 0.05 considered statistically significant.

## Acknowledgements

This project was supported by an innovation credit (IK17088) from the Ministry of Economic Affairs and Climate of the Netherlands.

This work was partly funded by the Dutch Cancer Society KWF 2022-14364 (to M.M.A.V).

## Author contributions

Author contributions: Conceptualization: KQ, JJ, PV, and HL. Methodology, patient biopsies and details: OMO, AR, MMAV and GvT. Image processing and analysis: TO, AO, OMO and ST. Automation: JH and AS. Data curation and analysis: OMO, CN, KQ, ST, AO and JH. Supervision: KQ and HLL. Writing-draft preparation: KQ and OMO. Writing-review and editing: OMO, AR, DK, CN, ST, TO, AO, AS, JH, GT, MMAV, SJT, JJ, PV, HL and KQ.

## Abbreviations used in this paper

HCC: Hepatocellular carcinoma
OoC: Organ-on-a-Chip
HMG-CoA reductase: 3-hydroxy-3-methylglutaryl-coenzyme A reductase
MASH: Metabolic dysfunction-associated steatohepatitis
TME: Tumor microenvironment
ECM: Extracellular matrix
SoC: Standard of Care
tSNE: t-distributed stochastic neighbor embedding
PCA: Principal component analysis
MMPs: Matrix metalloproteinase
TIMP: Tissue inhibitors of metalloproteinases
TGFβ: Transforming growth factor beta
ILs: Interleukins
IL6: Interleukin 6
CXCLs: chemokine (C-X-C motif) ligand
CCL: chemokine (C-C motif) ligand
DMEM: Dubelcco’s Modified Eagle Medium
EMEM: Eagle’s Minimal Essential Medium
MV2: Endothelial Cell Growth Medium MV2
EGCM2: Endothelial Cell Growth Medium-2
BSA: Bovine serum albumin
CAF: Cancer-associated fibroblasts
HUVEC: Human umbilical vein endothelial cells
DTC: Dissociated tissue cells

