## Supplemental Information for "Modeling Hepatocellular Carcinoma and its microenvironment on a chip Hepatocellular carcinoma PDChip"

**Suplemmental Information**

**Hepatocellular carcinoma PDChip**

Orsola Mocellin[1], Stephane Treillard[1], Abbie Robinson[1], Aleksandra Olczyk[1], Thomas Olivier[1], Chee P. Ng[1], Arthur Stok[1], Gilles van Tienderen[2], Monique M.A. Verstegen[2], Jeroen Heijmans[1], Dorota Kurek[1], Sebastian J. Trietsch[1], Henriëtte L. Lanz[1], Paul Vulto[1], Jos Joore[1] and Karla Queiroz[1,*]

[1]MIMETAS BV, De Limes 7, NL-2342DH Oegstgeest, The Netherlands

[2]Department of Surgery, Erasmus MC Transplant Institute, Erasmus MC-University Medical Center Rotterdam, NL-3015GD Rotterdam, The Netherlands

**Inventory of Supplemental Information**

**Supplementary Figure 1, is related to Figure 3**

**HCC1**

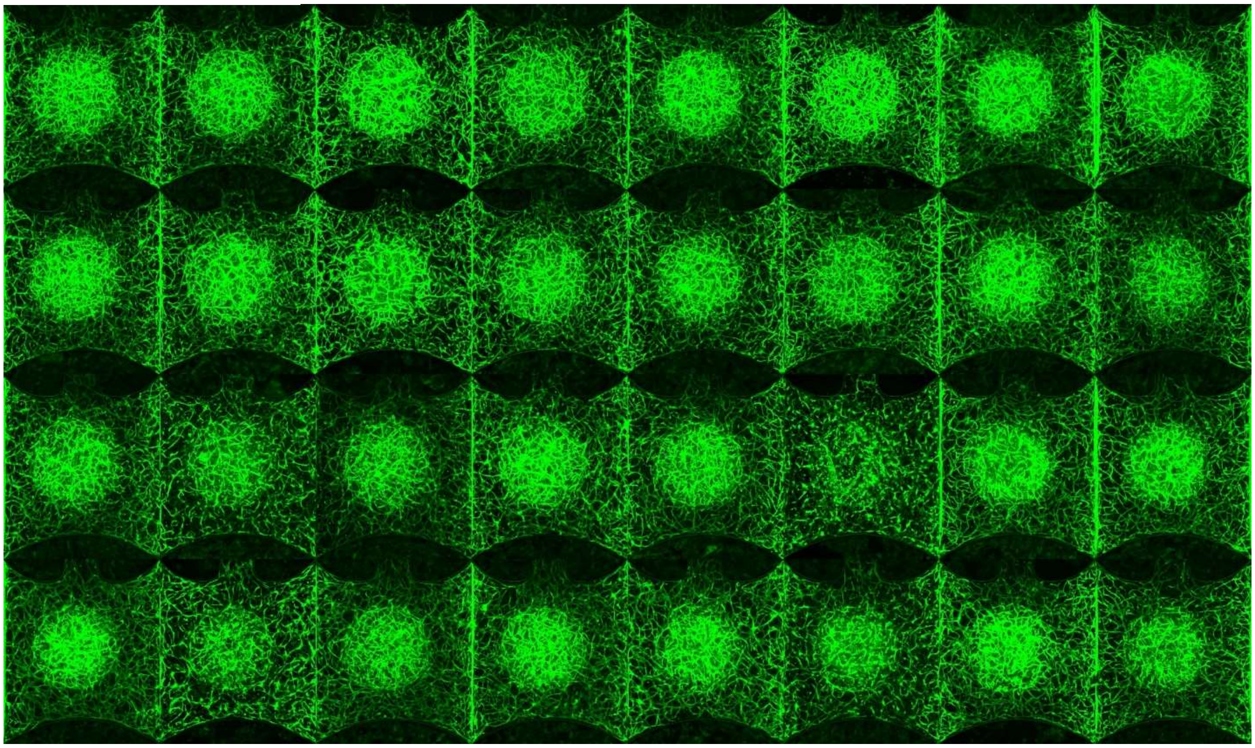

**HCC2**

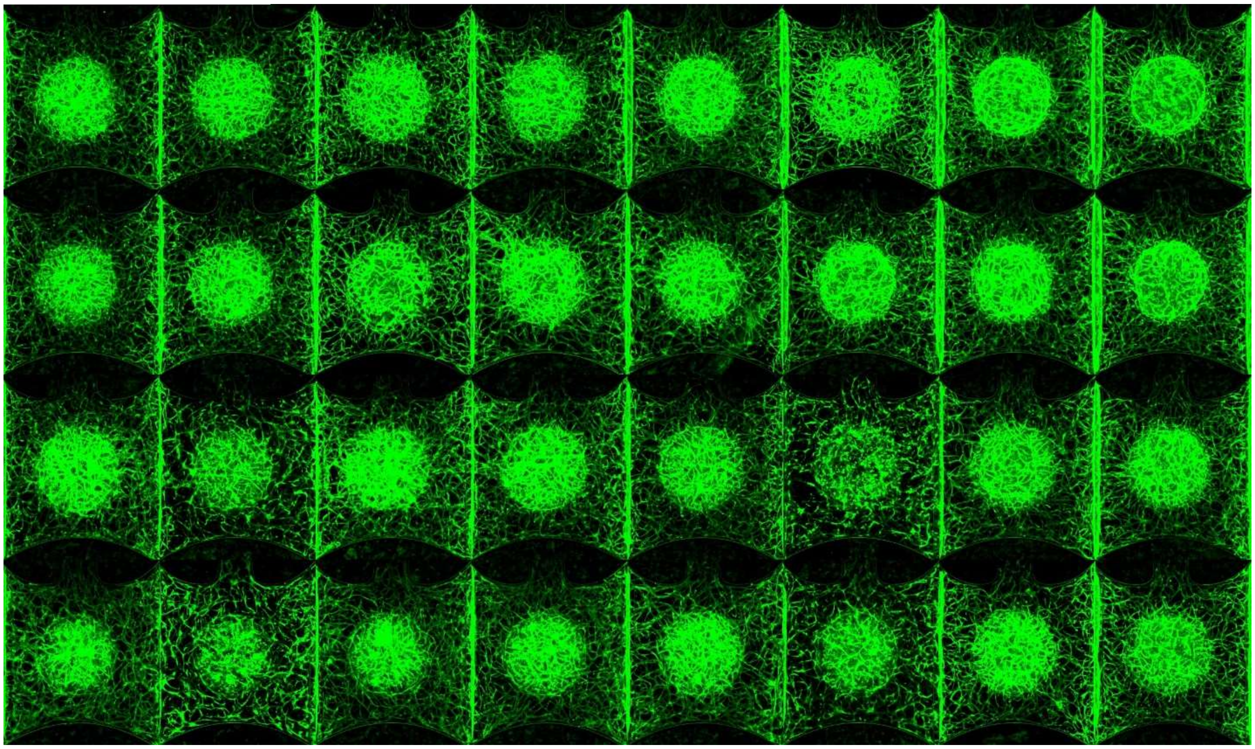

**HCC3**

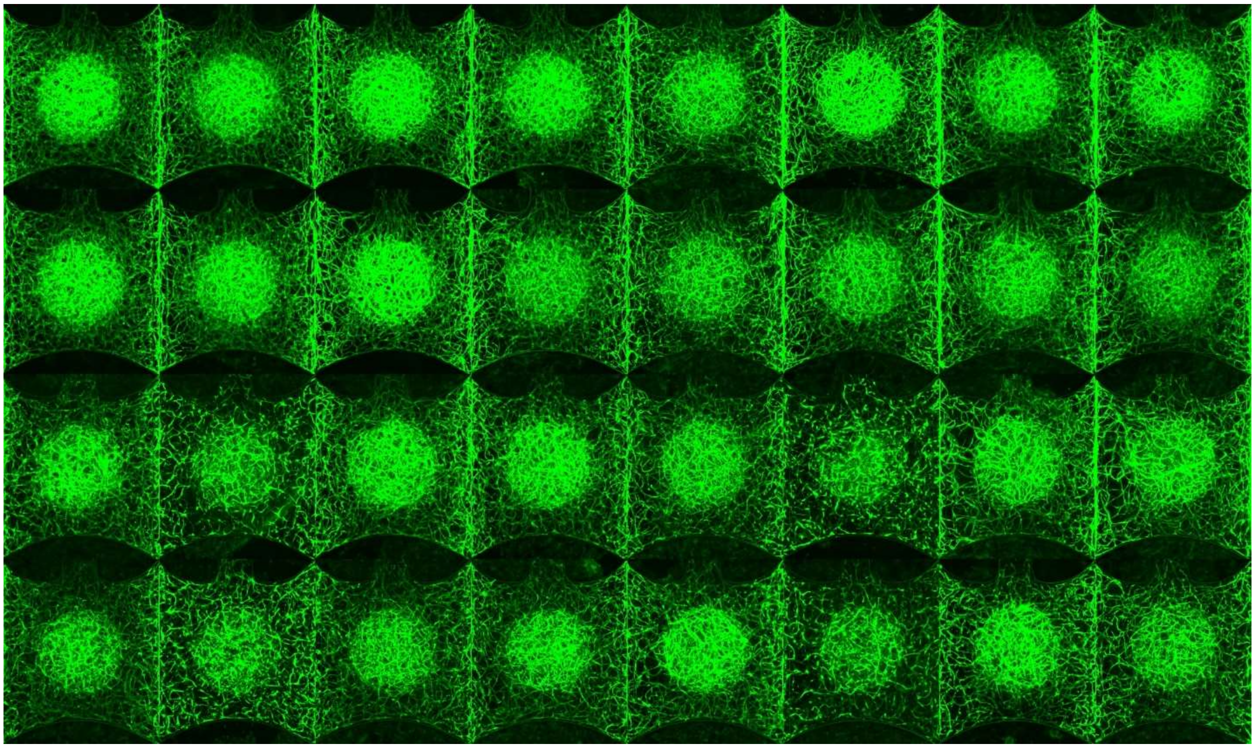

**HCC4**

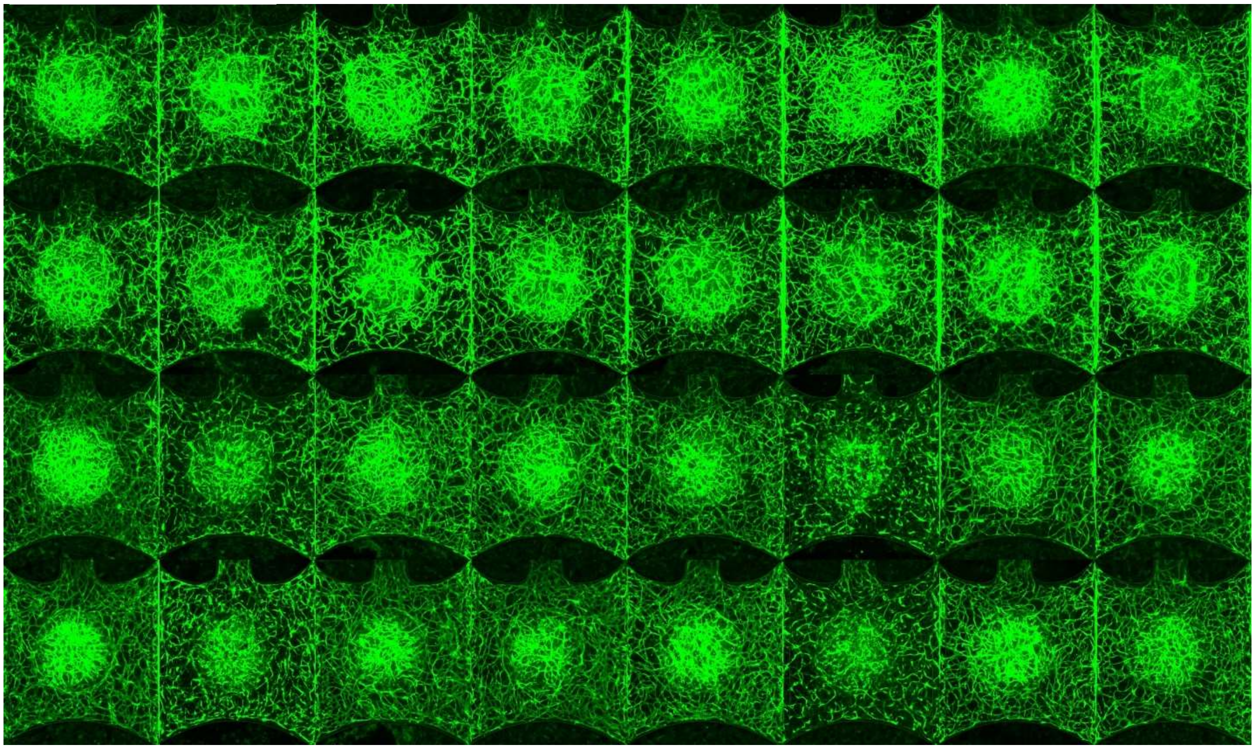

**HCC5**

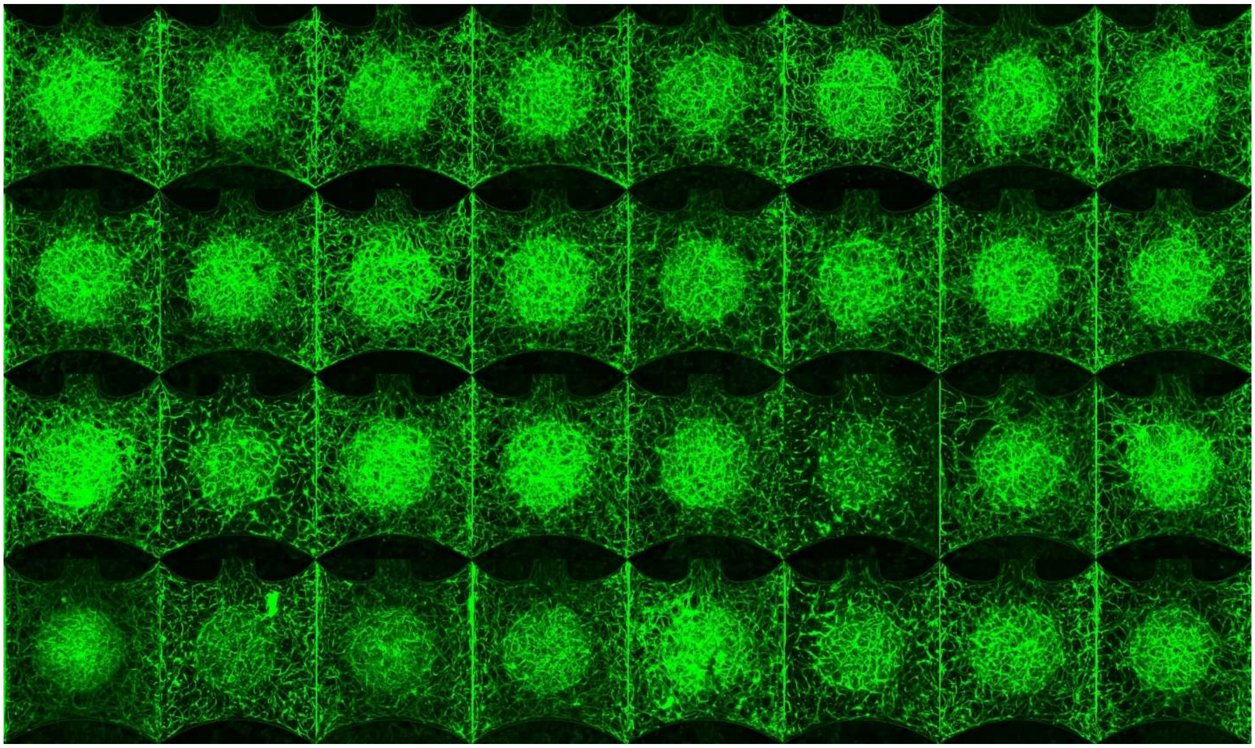

HCC6

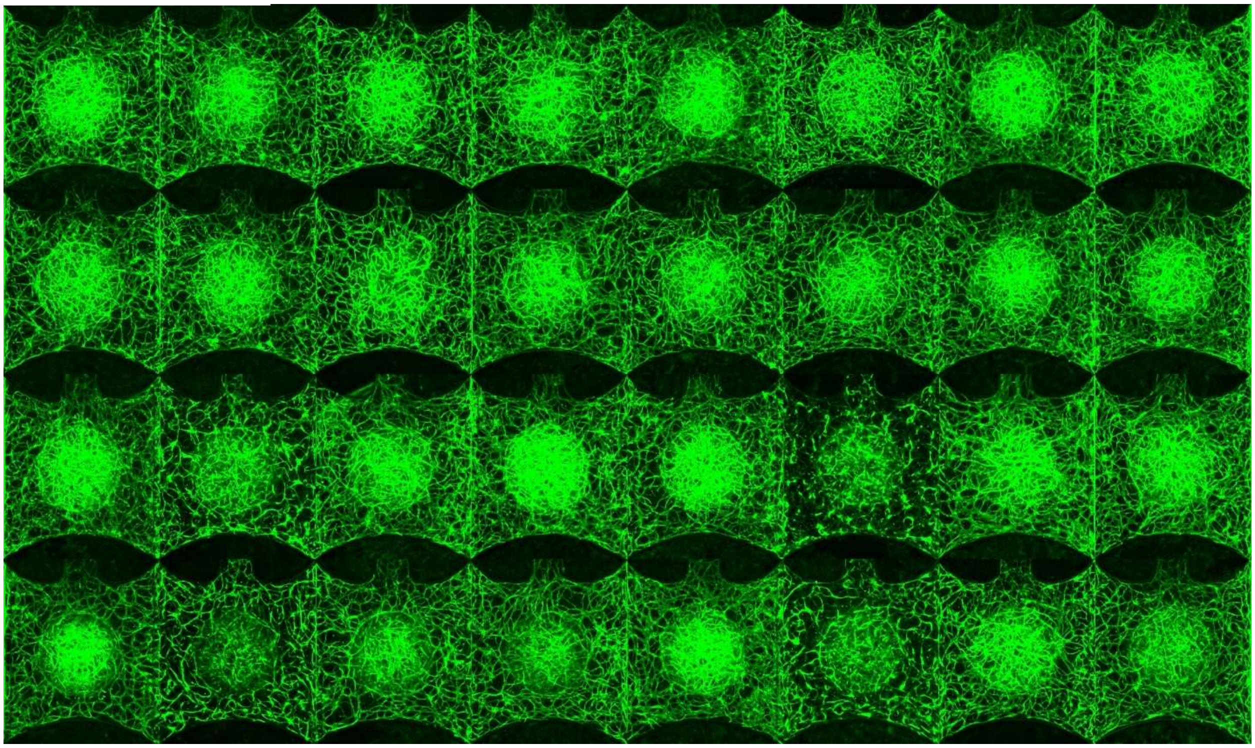

HCC7

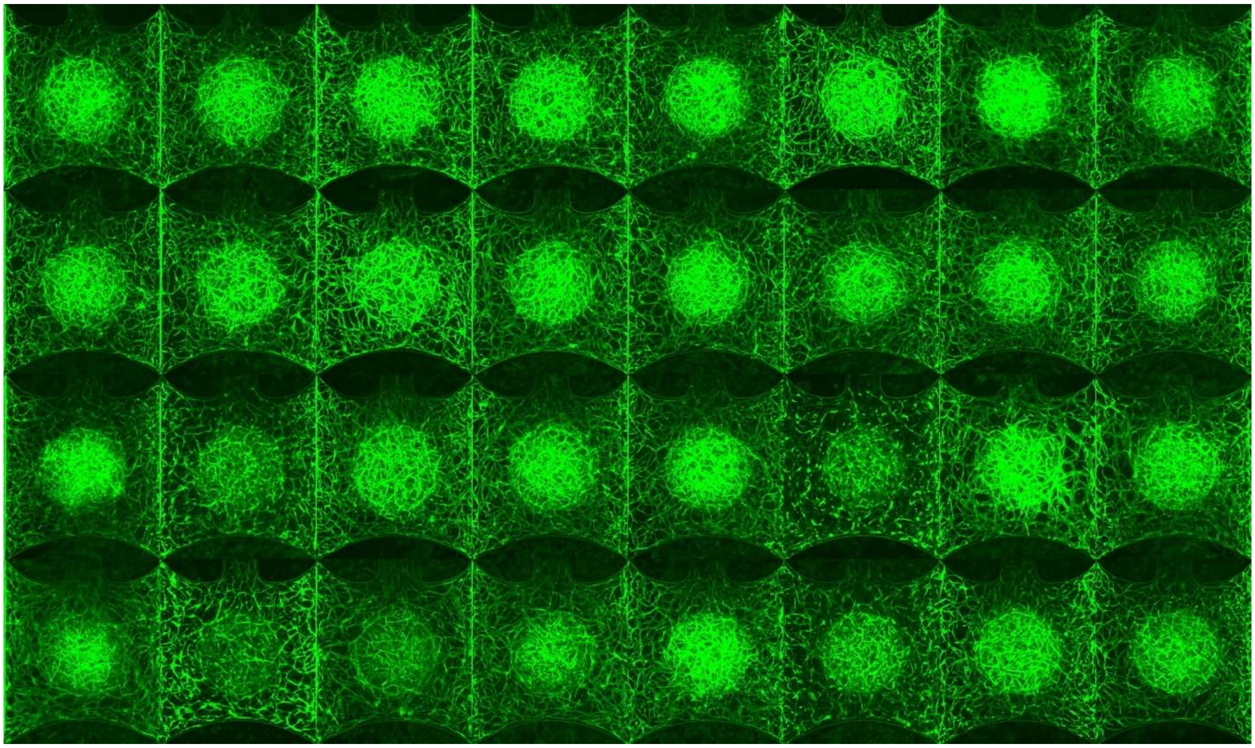

HCC8

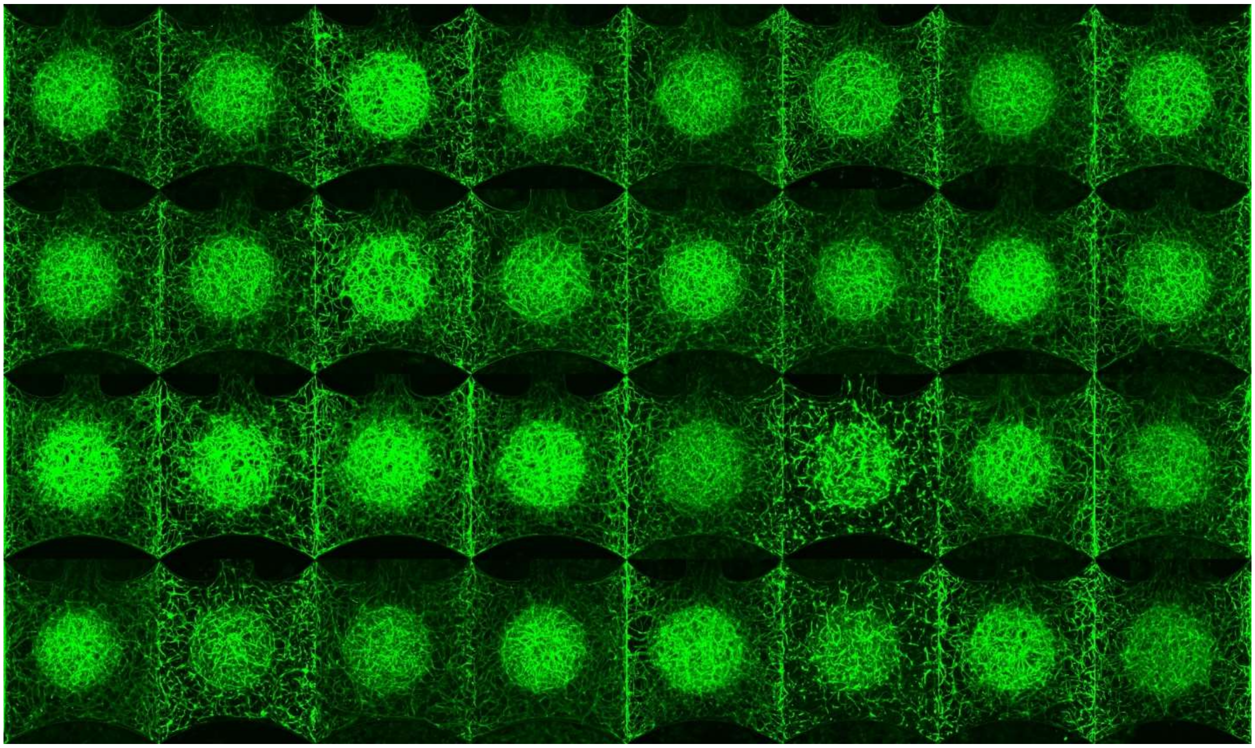

Huh7

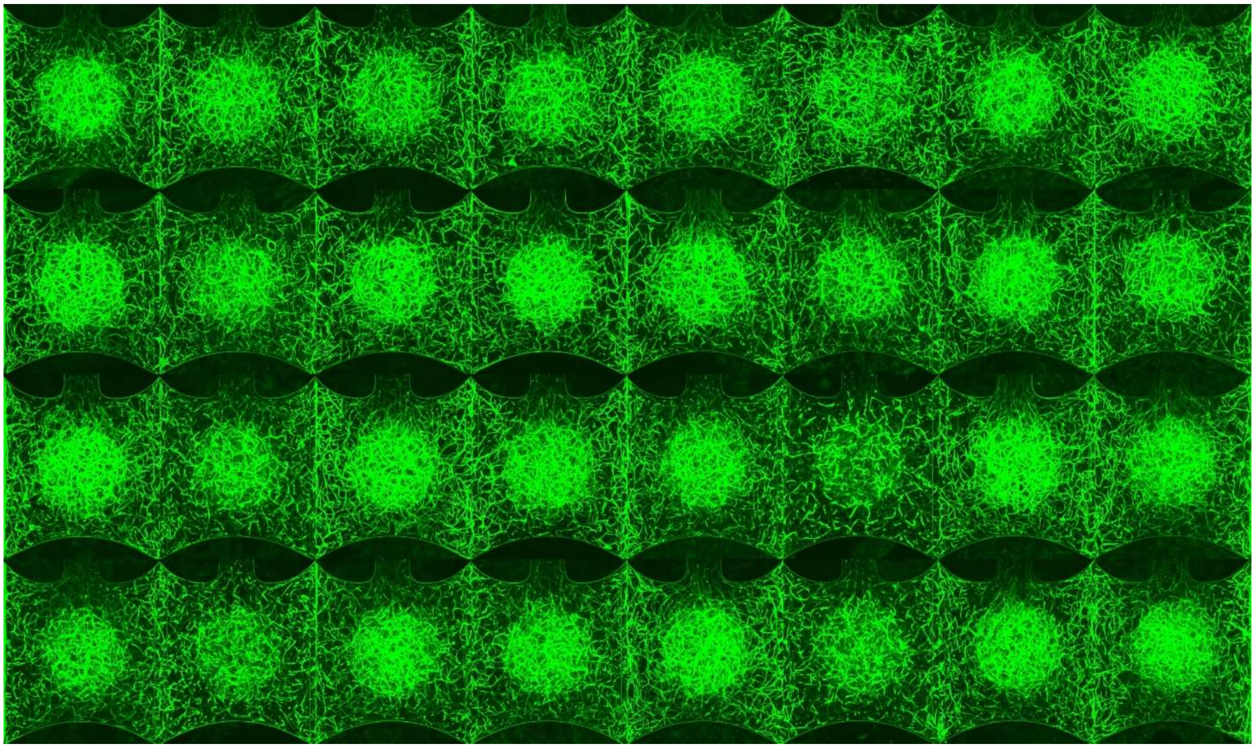

**HLE**

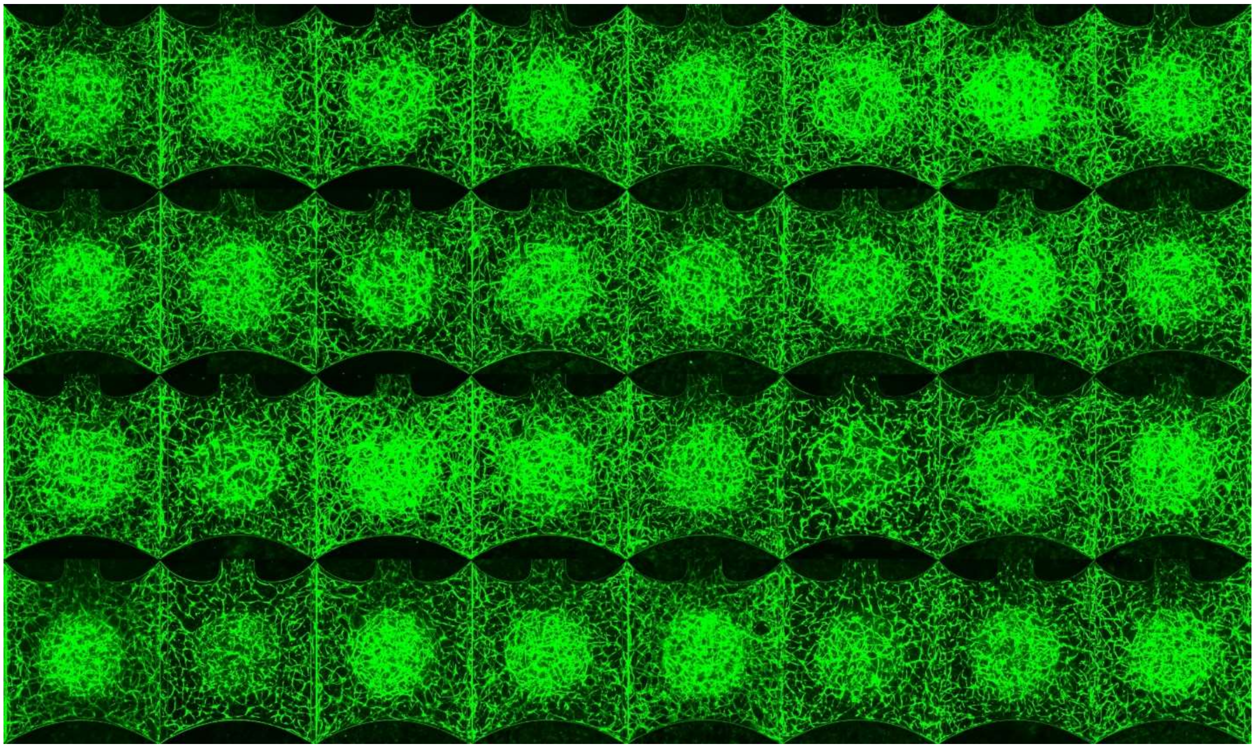

**Supplementary Figure 1. HCC PDChip-associated vasculature.** HCC vascularized constructs (HCC 1-8, Huh7 and HLE) were immunostained for CD31 (green). Immunostained HCC PDChips were confocal imaged and used for the phenotypical assessment of the vasculature.
